# Mechanosensitivity of nucleocytoplasmic transport

**DOI:** 10.1101/2021.07.23.453478

**Authors:** Ion Andreu, Ignasi Granero-Moya, Nimesh R. Chahare, Kessem Clein, Marc Molina Jordàn, Amy E. M. Beedle, Alberto Elosegui-Artola, Xavier Trepat, Barak Raveh, Pere Roca-Cusachs

## Abstract

Mechanical force controls fundamental cellular processes in health and disease, and increasing evidence shows that the nucleus both experiences and senses applied forces. Here we show that nuclear forces differentially control passive and facilitated nucleocytoplasmic transport, setting the rules for the mechanosensitivity of shuttling proteins. We demonstrate that nuclear force increases permeability across nuclear pore complexes, with a dependence on molecular weight that is stronger for passive than facilitated diffusion. Due to this differential effect, force leads to the translocation into or out of the nucleus of cargoes within a given range of molecular weight and affinity for nuclear transport receptors. Further, we show that the mechanosensitivity of several transcriptional regulators can be both explained by this mechanism, and engineered exogenously by introducing appropriate nuclear localization signals. Our work sets a novel framework to understand mechanically induced signalling, with potential general applicability across signalling pathways and pathophysiological scenarios.

**One sentence summary:** Force application to the nucleus leads to nuclear accumulation of proteins by differentially affecting passive versus facilitated nucleocytoplasmic transport.

## Main Text

Cells sense and respond to mechanical stimuli from their environment by a process known as mechanosensing, which drives important processes in health and disease (*1*–*3*). Growing evidence shows that the cell nucleus is directly submitted to force (*4*–*6*), and can act as a mechanosensor (*7*). Force applied to the nucleus (henceforth termed nuclear force for simplicity) can affect chromatin architecture (*8*), the accessibility of the transcription machinery (*9*), the conformation of nucleoskeletal proteins such as lamins (*10*), or cell contractility (*11, 12*). Further, forces transmitted to cells, and specifically nuclei, affect the nucleocytoplasmic localization of transcriptional regulators involved in different signalling pathways (*13*). As proposed for MRTF-A (*14, 15*), β-catenin (*16, 17*), or YAP (*18*–*20*), this can be due to a retention mechanism, in which force controls the localization of proteins by regulating their affinity for binding partners in the nucleus or cytoplasm. Alternatively, the force-sensitive step could be the nucleocytoplasmic translocation itself, as suggested also for YAP (*6*) and MyoD (*21*). This opens the hypothesis that nucleocytoplasmic transport could be mechanosensitive *per se*, independently of any specific signalling pathway. This would enable a general mechanism by which nuclear force could control the nuclear localization of proteins, and thereby transcription. However, whether there is such a mechanism, how it operates, and how it allows for directionality and molecular specificity, remains unknown.

Nucleocytoplasmic transport takes place through Nuclear Pore Complexes (NPCs) in two main ways, passive and facilitated diffusion (*22, 23*). Passive diffusion is rapid for small proteins, but is progressively impaired as the molecular weight (MW) of the protein increases (*24*–*26*). This impairment is caused by a meshwork of disordered proteins within NPCs called phenylalanine-glycine (FG) Nups, commonly termed the NPC permeability barrier (*27*). Facilitated diffusion of larger cargoes (proteins, RNA, and ribosomes) is mediated by various soluble nuclear transport receptors (NTRs) (*28, 29*). NTRs interact specifically with both the cargo molecules and FG Nups, thereby overcoming the NPC permeability barrier. They are divided between importins (mediating active nuclear import) and exportins (mediating active nuclear export) (*30*). Both classes interact with cargoes by binding to specific sequences (*31*) termed nuclear localisation signals (NLS, for proteins binding to importins) or nuclear export signals (NES, for proteins binding to exportins)(*32, 33*). The directionality of facilitated transport in either the import or export direction (for importins and exportins, respectively) is enabled by the coupling of binding/unbinding events to the phosphorylation status of the small GTPase Ran (either GTP, predominant in the nucleus, or GDP, predominant in the cytoplasm) (*29*). For example, in the so-called classical mode of import, an import complex is first formed between importin β (which interacts with FG nups), importin α (which binds importin β), and the cargo (which binds importin α through an NLS). The complex then diffuses through the NPC and finally dissociates in the nucleus in a RanGTP-dependent manner (*30, 31*).

To isolate how nuclear force affects nucleocytoplasmic transport, we studied different artificial constructs undergoing both passive and facilitated diffusion, but devoid of binding domains to partners in either the cytoplasm or nucleus. First, we used a light-inducible nuclear export construct (LEXY) (*34*) (Fig. 1A). Without excitation, the construct presents a mild NLS (cMyc^P1A^ NLS) fused to a mCherry and a folded LOV2 domain from *Avena sativa* phototropin-1 (AsLOV2). Under excitation with light (488 nm), the AsLOV2 domain unfolds to present a C-terminal encoded nuclear export signal (NES) that is stronger than the NLS. We transfected the construct in mouse embryonic fibroblasts (MEFs). To control the mechanical environment, cells were seeded on soft or stiff fibronectin-coated polyacrylamide gels (Young’s modulus of 1.5 and 30 kPa, respectively). Before photoactivation (t=0), with only the NLS active, the nuclear to cytoplasmic ratio (N/C ratio) was higher for cells on stiff substrates (Fig. 1B,C). Upon excitation by light, the construct exited the nucleus to similar final N/C ratios in both conditions, although the rate of N/C change (obtained by fitting an exponential equation to the curve) was higher for the stiff substrate (Fig. 1B-D). Once light excitation stopped, the reverse process occurred, with N/C ratios increasing faster for the stiff substrate, until restoring original values (Fig. 1E). We then co-transfected cells with DN-KASH, a dominant-negative domain of nesprin that prevents binding between nesprin and sun, two fundamental components of the Linker of Nucleus and Cytoskeleton (LINC) complex (*4*). By disrupting the LINC complex, DN-KASH has been shown to prevent force transmission to the nucleus on stiff substrates (*6*). DN-KASH overexpression led cells on stiff substrates to behave like those on soft substrates (Fig. 1B-E), demonstrating that the effect of stiffness was mediated by force transmission to the nucleus.

**Figure 1.**
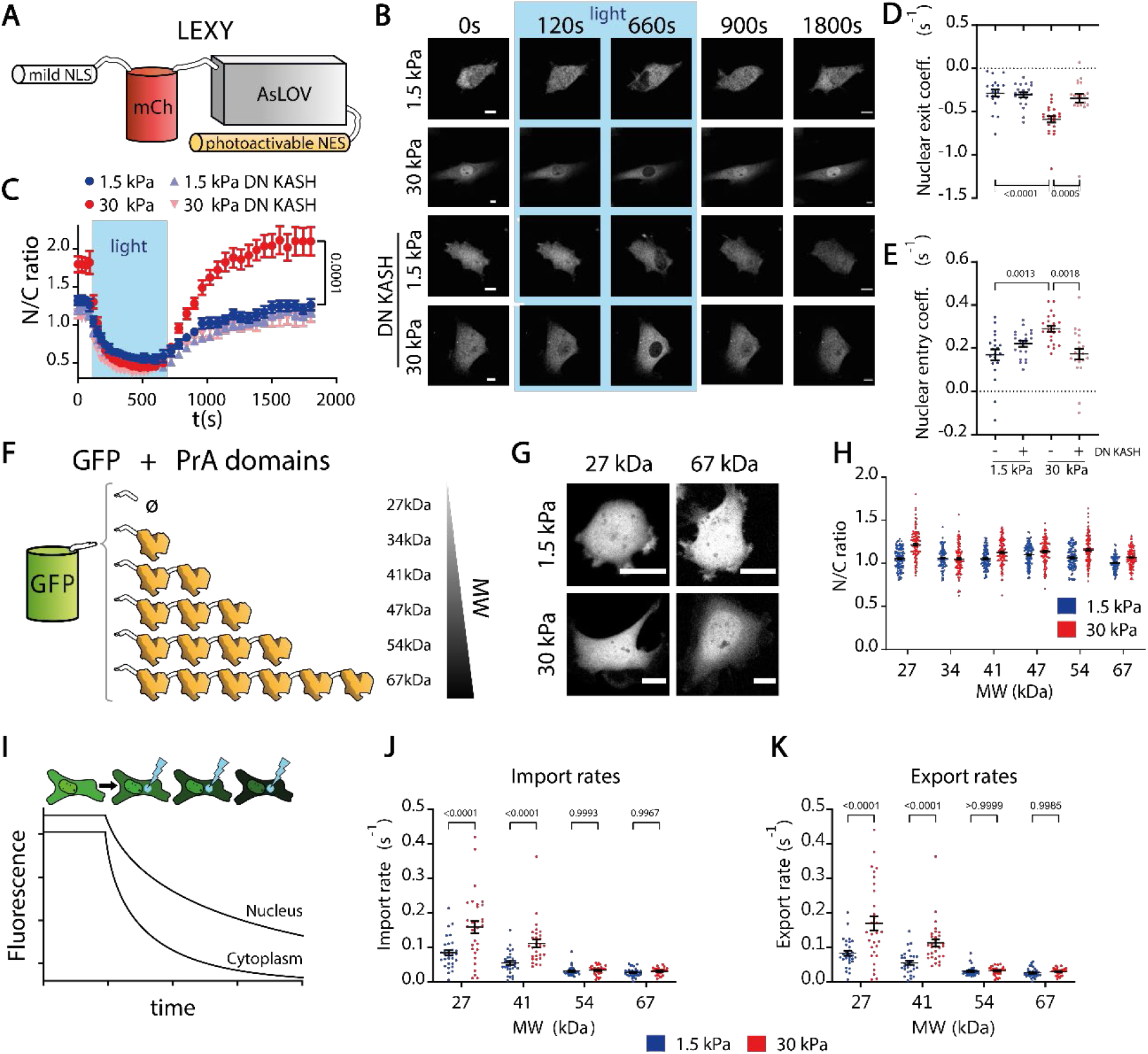
Nucleocytoplasmic transport is mechanosensitive. **A)** Cartoon of light-activated nucleocytoplasmic shuttling construct. Mild NLS is always active, NES is activated only upon light excitation. **B)** Time sequences of construct fluorescence before, during, and after excitation for cells seeded on 1.5/30 kPa substrates, with or without DN KASH overexpression. **C-E)** Corresponding quantifications of N/C ratios, and coefficients of exit and subsequent re-entry of constructs into the nucleus (in units of s^-1^, obtained by fitting an exponential to the curves, see methods). (N≥20 cells per condition from 3 independent experiments, p-values calculated with Mann-Whitney test; in C) the bar indicates the statistical significance between the last timepoint of 1.5kPa and 30kPa values). **F)** Cartoon of constructs with EGFP and different amount of repeats of PrA domains. **G)** Images showing fluorescence of indicated constructs on 1.5/30 kPa substrates. **H)** N/C ratios of constructs on 1.5/30 kPa substrates as a function of MW. N=120 cells from 3 independent experiments. Significant effects of stiffness and MW were observed (p <0,0001 and p <0,0001; computed via 2-way ANOVA). **I)** Cartoon depicting FLIP measurements: a laser photobleaches a region of the cell cytoplasm, and fluorescence intensities are recorded over time in nucleus and cytoplasm. Resulting curves are fitted to a kinetic model to obtain import and export rates (see methods). **J**,**K)** Import and export rates on 1.5 and 30 kPa substrates as a function of MW of the constructs. N=30 cells from 3 independent experiments. The effects of both substrate stiffness and MW were significant in both (J,K) cases (all p-values<0.0001). Scale bars, 20 µm. Data are mean ±S.E.M.

These results strongly suggest that nucleocytoplasmic transport is indeed generally affected by nuclear force, but do not distinguish between the contributions of passive and facilitated diffusion (since the ∼45 KDa LEXY construct is likely sufficiently small to diffuse passively). To dissect the different contributions, we first used constructs undergoing only purely passive diffusion, and regulated their diffusivity through changes in their MW. These constructs were composed of a Green Fluorescent Protein (GFP), attached through a short linker to between zero and six repeats of the 7 kDa bacterial Protein A (PrA) (Fig. 1F). PrA is inert and purely diffusive in eukaryotic cells, as shown previously (*24*) and also confirmed by the complete fluorescence recovery of the constructs after photobleaching either nucleus or cytoplasm (Fig. S1E). As such, these constructs have been previously used to study the progressive decrease in diffusion through NPCs with increasing molecular weight (MW) (*24*). When we transfected the constructs in cells, the N/C ratios of all proteins were ≈ 1 regardless of MW and substrate stiffness (Fig. 1 G,H).

This result shows that steady state concentrations of passively diffusing proteins were not mechanosensitive (where mechanosensitivity is defined throughout the manuscript as the fold change in a given magnitude in stiff versus soft substrates). However, this does not provide information on the effect of force on diffusion kinetics. To quantify this effect, we adapted a previously described method and model (*20*) based on Fluorescence Loss in Photobleaching (FLIP, Fig. 1I). Briefly, by progressively photobleaching the cell cytoplasm while simultaneously acquiring images, one can fit the resulting florescence decay curves in both cytoplasm and nucleus to an appropriate kinetic model, thereby obtaining nuclear import and export rates (see methods and Fig. S1). As expected, both import and export rates decreased with MW (Fig. 1J,K). Interestingly, rates increased with substrate stiffness, and this effect decreased for increasing MW (Fig. 1J,K). Confirming that this was mediated by nuclear force, DN-KASH overexpression had the same effect as reducing substrate stiffness (Fig. S2). Thus, nuclear force weakens the permeability barrier of NPCs (i.e., increases diffusion), and the effect is more pronounced for molecules with low MW (high diffusivity). Nevertheless, and because diffusion is non-directional, this does not affect the steady state nucleocytoplasmic distribution of molecules, which remains uniform.

Next, we assessed how nuclear force affected facilitated transport. To this end, we first assessed the behaviour of the protein directly interacting with FG nups, importin β, by transfecting cells with importin β-GFP. As expected by its affinity to FG nups, importin β-GFP localized at the nuclear membrane (Fig. 2A). Due to this localization and the diffraction limit, our FLIP measurements could not capture the likely very fast kinetics taking place in the immediate vicinity of the nuclear membrane. However, we did measure the kinetics of importin β molecules passing through nuclear pores and getting released in the bulk of either the nucleus or cytoplasm. Both import and export rates of importin β showed a high mechanosensitivity (Fig. 2B,C), similarly to that of highly diffusive passive molecules (Fig. 1J,K). Because importin β exhibits facilitated diffusion both in the import and export directions, import and export rates were largely symmetrical, leading to uniform concentrations inside and outside the nucleus regardless of substrate stiffness (Fig. 2D).

**Figure 2.**
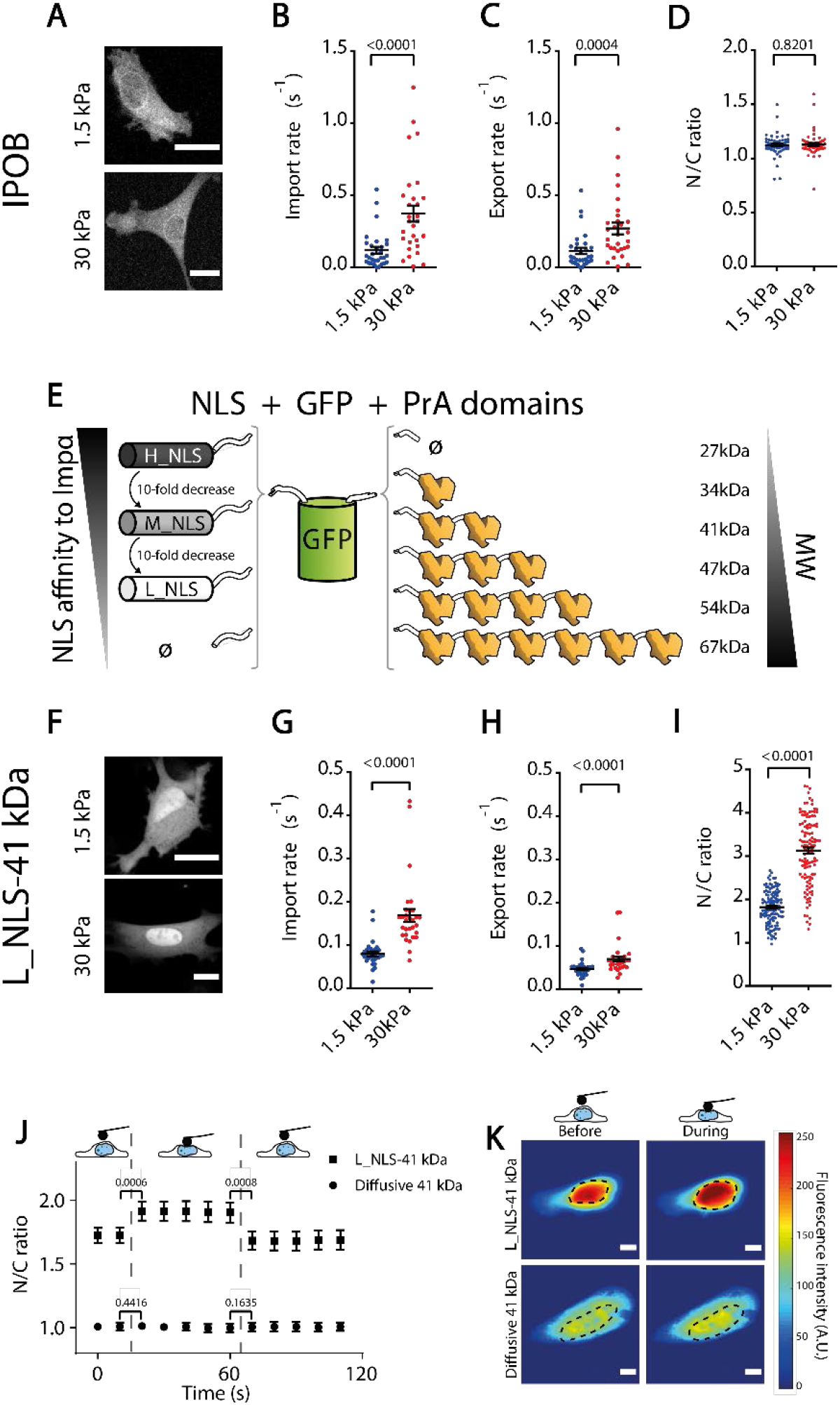
Higher mechanosensitivity of facilitated import versus passive diffusion explains force-induced nuclear translocation. **A)** Example importin β-GFP images for cells on 1.5/30 kPa substrates. **B-D)** Corresponding importin β-GFP import rates (B), export rates (C), and resulting N/C ratios (D). N=30, 30, and 60 cells from 3 independent experiments. p-values calculated with Mann-Whitney test. **E)** Cartoon of constructs with EGFP, different number of repeats of PrA domains, and NLS of different affinities to importin α. **F)** Example images of L_NLS-41 kDa construct for cells on 1.5 and 30 kPa substrates. **G-I)** Corresponding Import rates (G), export rates (H), and resulting N/C ratios (I) of L_NLS-41 kDa construct. N=30, N=30, N=120 cells from 3 independent experiments respectively each. p-values calculated with Mann-Whitney test. **I)** Corresponding images of L_NLS-41 kDa construct. **J)** N/C ratios of L_NLS-41 kDa or diffusive 41 kDa constructs in cells seeded on 1.5 kPa gels before, during, and after nuclear deformation with AFM. N= 16 cells from 3 independent experiments. p-values were calculated with a paired t-test. **K**) Corresponding images of constructs before and during force application, dotted line marks nucleus outline. Scale bars 20µm. Data are mean ±S.E.M.

Then, we studied the behaviour of cargo proteins undergoing facilitated diffusion. To this end, we added NLS sequences to the different GFP-PrA constructs (Fig. 2E). To regulate facilitated diffusion, we used different previously described functional NLS sequences with varying levels of affinity for importin α (*35*) (see table S2). The sequences ranged from that of the simian virus 40 (SV40), with very high affinity (which we termed H_NLS), to progressively lower affinities obtained by different point mutations in the sequence (which we termed M_NLS and L_NLS, for medium and low affinity). This approach allowed us to independently control passive and facilitated diffusion by regulating the number of PrA repeats and the NLS sequence, respectively. Interestingly, the mechanosensitivity of such constructs can be already predicted from the kinetic behaviour of passively diffusing molecules (Fig. 1G,H) and importin β (Fig. 2B,C), even if both showed uniform nucleo-cytoplasmic distributions regardless of stiffness. Indeed, a cargo molecule with an NLS should have a high mechanosensitivity in the import direction (because it enters the nucleus with importin β), but a low mechanosensitivity in the export direction if its MW is above ∼ 40 kDa (because it exits the nucleus through passive diffusion, which loses mechanosensitivity as MW increases).

By taking L_NLS-EGFP-2PrA (41 kDa) as a starting point, we confirmed this prediction: this molecule had a higher mechanosensitivity in import than export rates, leading to an increase in N/C ratios with stiffness (Fig. 2F-I). Confirming that this was mediated by nuclear force, the same effects on rates were observed when comparing cells with and without DN-KASH overexpression (Fig. S2). Further, the increase in N/C ratios was replicated by applying force to the nucleus of cells seeded on soft gels with an Atomic Force Microscope (AFM) (Fig. 2J,K). Interestingly, the fast decrease in N/C ratio upon force release in the AFM experiment shows that this mechanism is reversible in the timescale of seconds. As a control, force application with the AFM had no effect on the equivalent purely diffusive construct (Fig. 2J,K).

For the specific case of L_NLS-EGFP-2PrA, our results thus show that nuclear accumulation with force of NLS-cargo proteins is explained by a higher mechanosensitivity of facilitated versus passive diffusion. To understand this differential behaviour, we hipothesized that it may arise from the role of MW. Indeed, passive diffusion is strongly impaired as MW increases (*24*), whereas facilitated diffusion is expected to have a milder dependence on cargo MW (*36*–*38*). Thus, one could expect a scheme (summarized in fig. 3A) in which passive diffusion decreases both in magnitude and in mechanosensitivity as MW increases (as measured in fig. 1J,K) whereas facilitated transport is not affected (or only mildly affected) by MW. To verify this hypothesis, we measured import and export rates of constructs containing the L_NLS sequence and different MW (Fig. 3B,C). Indeed, import rates (dominated by active transport, fig. 3B) had a much milder dependence on MW than export rates (dominated by diffusion and with very similar behaviour to that of purely diffusive constructs, Fig. 3C).

**Figure 3.**
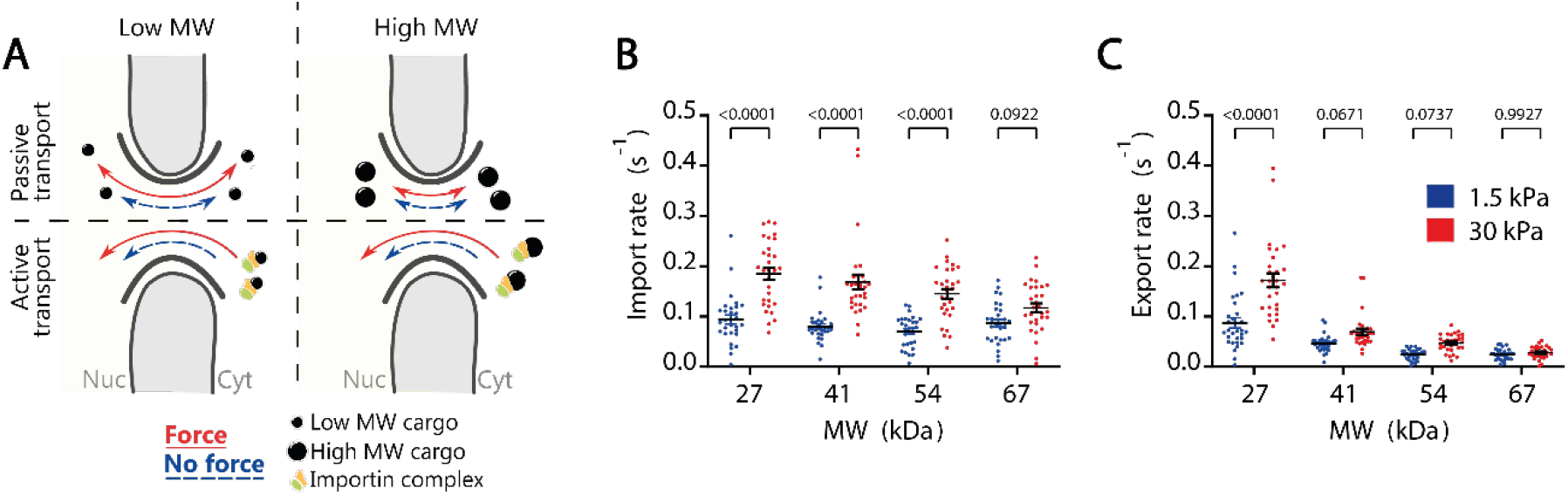
Forces to the nucleus increase facilitated and passive transport rates differentially. **A)** Cartoon summarizing the effects of nuclear force and MW on active and passive transport. Passive transport decreases with MW, and depends on force only for low MW molecules. Active transport does not depend on MW, and depends on force regardless of MW. **B)** Import rates (mediated by facilitated transport) of L_NLS constructs with different molecular weights. The effect of substrate stiffness and MW tested p<0.0001 and p=0.0004. **C)** Export rates of L_NLS constructs (mediated by passive transport) with different molecular weights. The effect of substrate stiffness and MW tested p<0.0001 and p<0.0001. N= 30 from 3 independent experiments. p-values from Two-way ANOVA. Data are mean ±S.E.M.

With these elements, we can generate a simple conceptual prediction of how nucleocytoplasmic transport should depend on force, MW, and NLS affinity. To this end, we assume that N/C ratios are given by the ratio of import and export rates, where export rates are purely passive and import rates have additive contributions of both passive and facilitated diffusion. Then, we assume as experimentally verified that **i)** passive import and export rates (which are equal) decrease as MW increases, **ii)** passive import and export rates increase when nuclear force is applied, but this effect disappears as MW increases, **iii)** facilitated import rates increase with nuclear force and with NLS sequence affinity, but do not depend on MW. We also assume that there is a limit to the efficiency of active facilitated transport, and therefore **iv)** N/C ratios saturate and cannot increase above a given level. With these assumptions, we can plot two simple diagrams showing how N/C ratios should depend on MW and NLS affinity before applying force to the nucleus (Fig. 4A), and their fold change with force, i.e., their mechanosensitivity (Fig. 4B). According to this framework, for low MW or a weak NLS, passive diffusion dominates over facilitated import, leading to N/C ratios close to 1 independently of nuclear force. For high MW or a strong NLS, facilitated import dominates over diffusion, leading to high, saturated N/C ratios, also independently of nuclear force. However, when passive and facilitated transport rates are comparable they depend differently on force, leading to mechanosensitive N/C ratios. As MW decreases (and passive diffusion increases) progressively higher facilitated import rates are required to match passive diffusion rates, and thus the “mechanosensitive zone” is placed along a diagonal in fig. 4B.

**Figure 4.**
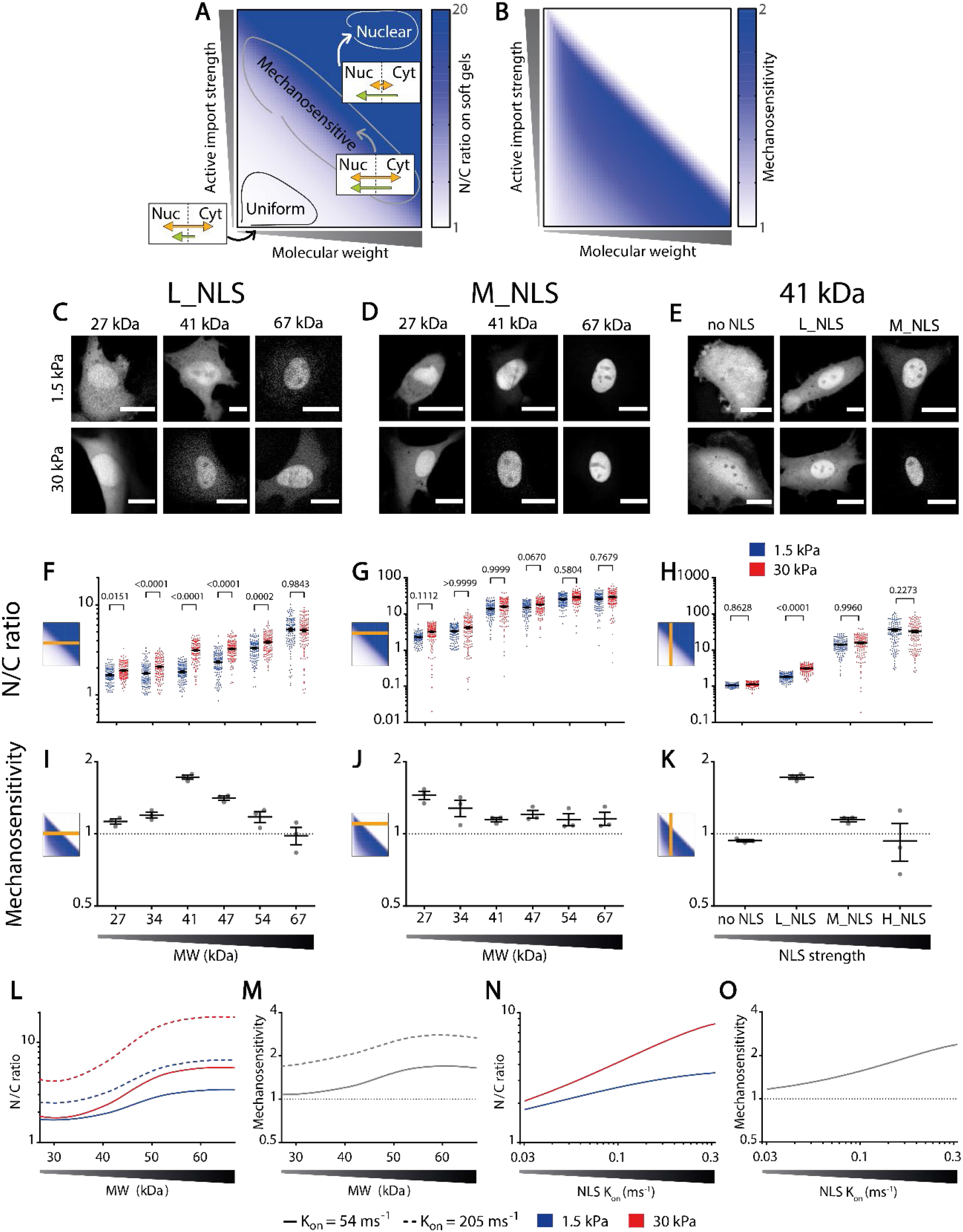
Balance between affinity to importins and MW defines the mechanosensitivity of nuclear localization. **A**,**B)** Qualitative prediction of how MW and affinity to importins should affect N/C ratios (A) on soft substrates and their mechanosensitivity (B) (see methods). Mechanosensitivity is defined as (N/C)_stiff_/(N/C)_soft_. **C-E)** Representative examples of construct distribution in cells seeded in substrates of 1.5kPa or 30kPa, for L_NLS constructs at different MW, M_NLS constructs at different MW, and 41kDa constructs at different NLS strengths. **F-H)** N/C ratios corresponding to the same conditions as C-E. **I-K)** Mechanosensitivity corresponding to the same conditions as C-E. **L-M)** Model predictions of N/C ratios (L) and mechanosensitivities (M) for NLS of different affinities for importin α (modelled through the binding rates k_on_ between the NLS and importin α, with values of 54 and 205 ms^-1^) as a function of MW. **N-O)** Model predictions of N/C ratios (N) and mechanosensitivities (O) for 41kDa constructs, as a function of increasing NLS strength. Statistics: All data were produced in 3 different repeats. F) N= 120 cells from 3 independent experiments. Both MW (p<0,0001) and Stiffness (p<0,0001) effects tested significant. G) N= 120 cells from 3 independent experiments. Both MW (p<0,0001) and Stiffness (p=0,0015) effects tested significant. H) N= 120 cells from 3 independent experiments. Both NLS strength (p<0,0001) and Stiffness (p=0,0012) effects tested significant. Adjusted p-values from 2-way ANOVA; Šídák’s multiple comparisons test. Scale bars: 20 µm. Data are mean ±S.E.M.

We then verified the different predictions by using the different constructs. First, increasing MW in proteins with a fixed NLS sequence (L_NLS) should progressively increase their N/C ratio, because the relative contribution of passive diffusion progressively decreases. Additionally, mechanosensitivity should be maximal at an intermediate range of MW between the high passive diffusion regime (low MW) and the saturated regime (high MW). Both trends were observed (Fig. 4C,F,I). Second, increasing MW in proteins with a fixed NLS sequence of higher affinity (M_NLS) should show the same trends found in low affinity NLS constructs. However, the point of maximum mechanosensitivity should happen at a lower MW, because the higher affinity NLS can more easily overcome the purely diffusive regime. This was also confirmed (Fig. 4D,G,J). Finally, increasing NLS affinity in proteins with a fixed MW (41 kDa) should progressively increase their N/C ratios, because the contribution of facilitated diffusion increases. In this case, mechanosensitivity should also be maximum at an intermediate range of NLS affinity, between the regime dominated by passive diffusion (low NLS affinity) and the saturated regime (high NLS affinity). This was also verified (Fig. 4E,H,K).

To test whether our experimental results could indeed be explained merely by changes in NPC permeability, in both passive and facilitated diffusion, we developed a more elaborate mathematical model that builds on the simple proposed conceptual framework but better accounts for the complexity of the nucleocytoplasmic transport cycle. The model describes transport kinetics using a system of ordinary differential equations, and as much as possible, it follows a canonical description of importin-mediated nucleocytoplasmic transport and the Ran cycle (see methods) (*30, 39*–*41*). To model the effect of force on passive diffusion, we used the experimentally measured passive diffusion rates as a function of force and MW from fig. 1J,K. For facilitated diffusion, we simply assumed that force reduces the mean time required for importin-cargo complexes to cross nuclear pores (in a MW-independent way), without changing any other parameter, thus maintaining a minimal number of free parameters. The model correctly predicted the increase of N/C ratios, and of their mechanosensitivity, with MW and NLS affinity (Fig. 4L-O). In contrast, the model did not capture the progressive saturation of N/C ratios at force-independent levels for high MWs or strong NLS sequences, and the associated decrease in mechanosensitivity. Instead, increasing NLS affinity to very high values in the model led to a collapse (rather than saturation) of both N/C ratios and mechanosensitivities (Fig. S3). However, this model prediction actually explains the somewhat surprising experimental observation that export rates (and not only import rates) also increase with increasing NLS affinity (Fig. S3). According to the model, this is explained by very strongly bound cargo exiting the nucleus before unbinding from importins in the nucleus (Fig. S3). The model may overestimate this effect, which might explain the discrepancy between model and experiments at high MWs and NLS strengths (Fig. 4L-O; Fig. S3). Independently of the discrepancy between saturation and collapse in model and experiment, this effect on export rates can explain the loss of mechanosensitivity at very high N/C ratios: if export rates are mediated by facilitated rather than passive diffusion, then their dependency on force is the same as that of import rates, and the overall effect on N/C ratios cancels out.

Given the observed mechanosensitivity of active nuclear import, one might expect a similar (but reversed) behaviour for active export. To test this, we developed constructs by combining PrA repeats with different NES signals of different strength (*42*) (see table S2). N/C ratios changed as expected with MW and NES strength (by following the opposite trends than NLS constructs, fig. S4A-I). The mechanosensitivity of the constructs also behaved in the opposite way, with constructs leaving (rather than entering) the nucleus with force (Fig. S4G-I). Consistently, import and export rates of NES constructs also had opposite trends with MW than NLS constructs: export rates were largely independent of MW, whereas import rates showed a strong dependence, mimicking diffusive constructs (Fig. S4J,K). Interestingly, mechanosensitivity of the NES constructs was systematically milder than that of the NLS constructs. This is consistent with the behaviour of the light inducible construct (Fig. 1B), which had a stiffness-dependent localization when controlled by active import (no light excitation) but not when controlled by active export (under light excitation). This lower mechanosensitivity of active export as compared to import may be related to the many differences between the transport cycles in both directions (*30*). However, one potential intuitive explanation could be simply that generating a concentration gradient is more easily accomplished by accumulating proteins in a small compartment (the nucleus) than a large one (the cytoplasm). In line with this hypothesis, model predictions obtained by inverting nuclear and cytoplasmic volumes led to lower N/C ratios and mechanosensitivity (Fig. S4L-O).

Finally, we evaluated whether nucleocytoplasmic transport can explain the reported mechanosensitivity of different transcriptional regulators. Recent work has identified several transcriptional regulators which localize to the nucleus with force in different contexts, including YAP (*6, 43*), twist1 (*44*), snail (*45*), SMAD3 (*46*), GATA2 (*47*), and NFκβ (*48*). If their mechanosensitivity is explained by regulation of nucleocytoplasmic transport with nuclear force, then it should be abolished by preventing either force transmission to the nucleus (by overexpressing DN-KASH) or nucleocytoplasmic transport (by overexpressing either DN-Ran, a dominant-negative version of Ran (*49*), or by treatment with importazole, a drug which blocks active import by importin β (*50*)). For the case of YAP, we previously showed that its mechanosensitivity is abrogated by both factors (*6*). Regarding the rest, GATA2 and NFκβ exhibited a very low mechanosensitivity in our system (Fig. S5G,K), but SMAD3, Snail, and Twist1 showed a clear response (Fig. S5A-F and Fig. 5A,B). In all cases, mechanosensitivity was abrogated by DN-KASH, DN-RAN, or importazole (Fig. S5A-F and Fig. 5A,B). Interestingly and consistent with our finding that NLS constructs were more mechanosensitive than NES constructs, SMAD3 mechanosensitivity was higher for cells treated with TGFβ (which induces SMAD3 nuclear import) than with lapatinib (which induces SMAD3 nuclear export) (*51*).

**Figure 5.**
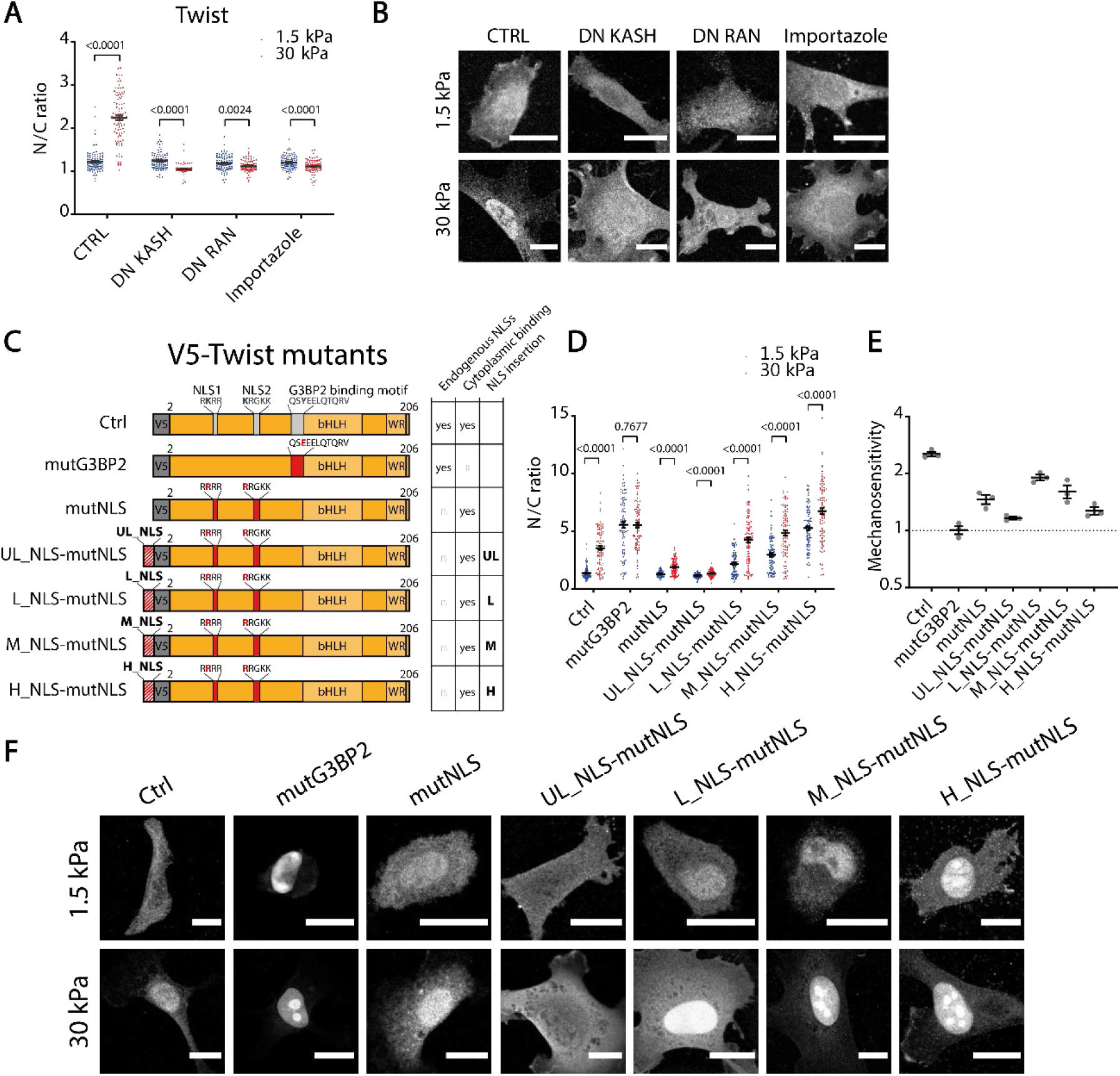
The mechanosensitivity of twist1 can be re-engineered with exogenous NLS sequences. **A)** N/C ratios of endogenous twist1 for cells on 1.5/30 kPa substrates, and under indicated treatments. N= 100 cells from 3 independent experiments. p-values from Mann-Whitney tests, corrected for multiple tests in the intracondition comparisons. **B)** Corresponding images of twist1 distribution. **C)** Scheme of different twist1 mutants. Mutations inactivating both NLS sequences and the G3BP2 binding motif are indicated in red. **D)** N/C ratios of transfected twist1 mutants for cells on 1.5/30 kPa substrates. N= 90 cells from 3 independent experiments. p-values from Mann-Whitney tests, corrected for multiple tests. **E)** Corresponding construct mechanosensitivities, defined as (N/C)_stiff_/(N/C)_soft_ (N= 3 experiments). **F)** Corresponding images showing the distribution of the different mutants. Scale bars, 20 µm, data are mean ±S.E.M.

Thus, the mechanosensitivity of several transcriptional regulators is indeed controlled by force-induced effects in nucleocytoplasmic transport. Our proposed mechanism also has the stronger implication that mechanosensitivity can be engineered simply by selecting the appropriate levels of affinity to importins. To verify this, we took twist1 as a convenient model, since its NLS sequences are well described, and their function can be abolished with simple point mutations (*52*). Further, its mechanosensitivity depends on its binding to G3BP2, which retains twist1 in the cytoplasm (*44*). We first overexpressed wild-type twist1 in cells, which retained the mechanosensitivity of endogenous twist1 (Fig. 5C-F). Then, we overexpressed a G3BP2 binding deficient mutant, mutG3BP2. As expected, this led to high N/C ratios on both soft and stiff substrates, thereby losing mechanosensitivity. Confirming the role of nucleocytoplasmic transport, the NLS dead mutant (mutNLS, still under the control of G3BP2), lost the nuclear localization in both soft and stiff substrates, thereby also losing mechanosensitivity (Fig. 5C-F). We then assessed whether we could restore twist1 mechanosensitivity by rescuing twist mutNLS not with its endogenous NLS, but by exogenously adding our different characterized NLS sequences (plus an additional ultra-low affinity sequence, UL_NLS). Indeed, adding NLS sequences of different strength mimicked the effects seen in fig. 4: as the NLS strength increased, nuclear localization progressively increased, and mechanosensitivity was highest at a low strength (L_NLS), where it was almost as high as in the endogenous case. Thus, simply substituting the endogenous twist1 NLS with an exogenous one of the appropriate strength, not regulated by any twist-1 related signalling mechanism, recapitulates its mechanosensitivity.

Our work shows that force regulates nucleocytoplasmic transport by weakening the permeability barrier of NPCs, affecting both passive and facilitated diffusion. Because MW affects more passive than facilitated diffusion, this generates a differential effect on both types of transport that enables force-induced nuclear (or cytosolic) localization of cargo. Three important open questions emerge from our findings. First, although NPCs have been reported to be flexible and thus potentially deformable under force (*53*–*55*), and their apparent size is bigger in nuclei under force (*6*), the specific structural changes in NPC structure induced by force, and how they weaken the permeability barrier, remain to be understood. In this regard, a recent preprint showed that the NPC central channel can constrict upon energy depletion in cells, which is likely related with a decrease in mechanical tension in the nucleus (*56*). Second, the exact set of properties that confer mechanosensitivity to transcriptional regulators or other proteins remains to be fully explored. The different transcriptional regulators discussed here range in size from over 20 kDa (for twist) to over 60 kDa (for YAP), thereby encompassing almost the full range of weights analyzed with our designed constructs. However, diffusivity through NPCs depends not only on MW but also on surface charges (*57*) and protein mechanical properties (*58*), which could play major roles. Finally, why facilitated export is less affected than facilitated import may be related to the different volumes of nucleus and cytoplasm (as suggested by modelling in fig. S4), to the different interactions between importins and exportins with FG-nups (*59*) or to the asymmetric manner in which NPCs deform (*56*), but is also unclear. Beyond these questions, our work demonstrates a general mechanism of mechanosensitivity, with incorporated specificity through molecular properties such as the NLS sequence and MW. Although other mechanisms (such as differential binding to nuclear or cytosolic proteins) can generate mechanosensitive nuclear translocation (*14, 16*), our mechanism is consistent with the behaviour of several transcriptional regulators, and has potential general applicability. Our findings suggest that interfering with nucleocytoplasmic transport may be an avenue to regulate or abrogate mechanically-induced transcription in several pathological conditions. Perhaps even more excitingly, they open the door to design artificial mechanosensitive transcription factors, to enable mechanical control of transcriptional programs at will.

## Methods

### Cell culture and reagents

Mouse embryonic fibroblasts (MEFs) were cultured as previously described (*60*), using Dulbecco’s modified eagle medium (DMEM, Thermofischer Scientific, 41965-039) supplemented with 10% FBS (Thermofischer Scientific, 10270-106), 1% penicillin-streptomycin (Thermofischer Scientific, 10378-016), and 1.5% HEPES 1M (Sigma Aldrich, H0887). Cell cultures were routinely checked for mycoplasma. CO2-independent media was prepared by using CO2-independent DMEM (Thermofischer Scientific, 18045-054) supplemented with 10% FBS, 1% penicillin-streptomycin, 1.5% HEPES 1M, and 2% L-Glutamine (Thermofischer Scientific, 25030-024). Media for AFM experiments was supplemented with Rutin (ThermoFischer Scientific, 132391000) 10 mg/l right before the experiment. Importazole (Sigma Aldrich) was used at 40 μM concentration for 1 h (*61*). Cells were transfected the day before the experiment using Neon transfection device (ThermoFischer Scientific) according to manufacturer’s instructions. Cells were seeded ∼4 h before the experiment.

### Antibodies and compounds

For primary antibodies, we used Anti Twist antibody (Twist2C1A, Santa cruz, sc-81417) 1:200, Mouse monoclonal antibody to SNAIL + SLUG - N-terminal (CL3700, abcam, ab224731) 1:200, rabbit polyclonal anti SMAD3 (Cell Signaling, 9513) 1:40, Rabbit polyclonal antibody to GATA2 (Abcam, ab153820) 1:200, rabbit polyclonal Anti-NF-kB p65 antibody (abcam, ab16502) 1:200. The secondary antibodies used were Alexa Fluor 488 anti-mouse (A-11029; Thermo Fischer Scientific) and Alexa Fluor 555 anti-rabbit (A-21429; Thermo Fischer Scientific), diluted 1:200.

### Plasmids

If not referred otherwise, plasmids were constructed via standard molecular biology methods. **LEXY plasmids:** NLS-mCherry-LEXY (pDN122) was a gift from Barbara Di Ventura & Roland Eils (Addgene plasmid # 72655; http://n2t.net/addgene:72655; RRID:Addgene_72655) (*34*). **Nuclear transport plasmids:** NLS, NES, or nought combinations with different molecular weight modules were designed as following: Localization signal plus GGGGS linker, EGFP, and different amount of Protein A from *Staphylococcus aureus* modules. Nuclear Localization Signals sequences were extracted from Hodel *et al*. (2001)(*35*). Nuclear Export Signals were extracted from Kanwal *et al*. (2004)(*42*). Protein A domain sequences were used originally in Timney *et al*. (2016) (*24*) and were kindly provided to us M. Rout. For more detailed information see tables S1 and S2. **DN-KASH DN-RAN:** DN (Dominant negative)-KASH was described previously as EGFP-Nesprin1-KASH in Zhang *et al*., (2001) (*62*). DN (Dominant negative)-RAN (Addgene plasmid # 30309, described as pmCherry-C1-RanQ69L) was a gift from Jay Brenman (*63*). **Twist mutants:** pBABE-puro-mTwist was a gift from Bob Weinberg (Addgene plasmid # 1783; http://n2t.net/addgene:1783; RRID:Addgene_1783) (*64*). mTwist was cloned into a pEGFP-C3 backbone and a V5 tag was included at the N-terminal. The different mutants were constructed by adding the corresponding NLS sequences and/or changing the indicated codons. For more detailed information see table S1.

### Polyacrylamyde gels

Polyacrylamide gels were prepared as previously described (*65*), and coated using a protocol adapted from the literature (*66*). Briefly, gels were covered with a mix containing 10% HEPES 0.5M Ph 6, 0.002% BisAcrylamide (BioRad), 0.3% 10 mg/ml N-hydroxysuccinimide (NHS, Sigma Aldrich) in dimethyl sulfoxide (DMSO, Sigma Aldrich), 1% Irgacure 2959 (BASF), 0,0012% Di(trimethylolpropane)tetra-acrylate (Sigma Aldrich), in milliQ water. Different concentrations of acrylamide and bis-acrylamide were used to obtain different stiffness gels. For 1.5 kPa gels we used 5,5% acrylamide and 0,04% bis-acrylamide; for 30 kPa we used 12% acrylamide and 0,15% bis-acrylamide. Gels were then illuminated with UV light for 10 minutes. After exposure, gels were washed once with HEPES 25mM Ph 6 and once with PBS. Gels were then incubated with 10 μg/ml of fibronectin in PBS overnight at 4ºC, UV treated in the hood for 10 minutes, once with PBS and immediately used. The rigidity of the gels was measured using Atomic Force Microscopy as previously described (*67*).

### Immunostaining

Immunostainings were performed as previously described (*6*). Cells were fixed with 4% paraformaldehyde for 10 minutes, permeabilized with 0.1% Triton X-100 for 40 minutes, blocked with 2% Fish-Gelatin in PBS 1X for 40 minutes, incubated with primary antibody for 1 hour, washed 3X with Fish-Gelatin-PBS for 5 minutes, incubated with secondary antibody for 1 hour, washed with Fish-Gelatin-PBS 3X for 5 minutes, and mounted using ProLong Gold Antifade Mountant (ThermoFischer Scientific).

### Steady state image acquisition and analysis

Cells were imaged with a Nikon Eclipse Ti inverted confocal microscope using a 60x water immersion objective 1.2 NA. Nuclear to cytoplasmic (N/C) ratios were quantified manually by segmenting the nucleus using Hoechst (immunostaining) or taking advantage of the GFP tagged construct (live cells) by the following formula:

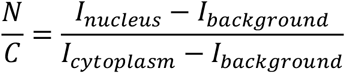

Where *I*_*nucleus*_ and *I*_*cytoplasm*_ are the mean fluorescence intensity of the nucleus and the cytoplasm respectively. ROIs in the nucleus an in the cytoplasm were selected manually next to each other, close to the nuclear membrane. *I*_*background*_ is the mean intensity of the background far from the cell.

Mechanosensitivity was calculated once for each of the repeats using the following formula:

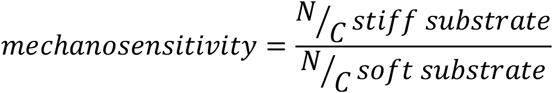

### Live cell AFM experiments

Live cell AFM experiments were carried out as previously described (*6*). Briefly, AFM experiments were carried out in a Nanowizard 4 AFM (JPK) mounted on top of a Nikon Ti Eclipse microscope. Polystyrene beads of 20 μm were attached using a non-fluorescent adhesive (NOA63, Norland Products) to the end of tipless MLCT cantilevers (Veeco). The spring constant of the cantilevers was calibrated by thermal tuning using the simple harmonic oscillator model. Experiments were carried out on cells previously transfected with the EGFP construct and incubated with Hoechst 33342 (Invitrogen), and seeded on gels on compliant gels. For each cell, the nucleus was identified by using the Hoechst fluorescence signal, and a force of 1.5 nN was applied to the nucleus. Once the maximum force was reached, the indentation was kept constant under force control, adjusting the z height by feedback control. An image was acquired every 10s by an Orca ER camera (Hamamatsu) and a 60X (NA = 1.2) objective.

### Photoactivation experiment and quantification

Photoactivation experiments were done with a Zeiss LSM880 inverted confocal microscope using a 63X 1.46 NA oil immersion objective. An argon laser was used with 561 nm wavelength for acquisition and 488 nm laser for stimulation. For the experiment 4 images were obtained before stimulation, then followed by 19 images during stimulation and 18 images for recovery. All images were acquired every 30 s and during the stimulation period the 488 nm laser was irradiated to the whole field of view during 1 s at 100% laser power.

To obtain the entry and exit coefficient a single exponential equation was fitted to the N/C ratio of each cell:

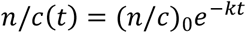

Where (*n*/*c*)_0_ is the initial ratio of the stimulation or recovery phase, *t* is time, and *k* is the entry or exit coefficient. The curve was fitted to the whole stimulation or recovery phase.

### FRAP Data Acquisition and Analysis

Estimation of mobile fraction of proteins was done using fluorescence recovery after photobleaching (FRAP) experiments. FRAP involves bleaching a region of interest (ROI) and then tracing recovery of fluorescence in that region with respect to time. Image acquisition was done with a Zeiss LSM880 inverted confocal microscope using a 63X 1.46 NA oil immersion objective and a 488nm wavelength argon laser at 100% lase power. We acquired images every 60 ms. We bleached and acquired images acquired every 60 ms for 12s. We use two ROIs for our experiments: first is the circular 14-pixel diameter (∼6.9 μm^2^) region being bleached (ROIF), and second is the cell area segmented manually (ROIC). The data for ROIs consist of the fluorescence integrated density as a function of time from images acquired before and after photobleaching. For further analysis, we normalize the fluorescence intensities of ROIs using the double normalization method (*68*). Double normalization corrects for photobleaching during the post bleach imaging and normalizes recovery fluorescence with a pre bleach signal. Double normalized intensity (I) for recovery signal can be calculated by using following formula.

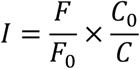

where *F* and *C* are the fluorescence integrated densities of ROIF and ROIC respectively for post bleach imaging, and *F*_*0*_ and *C*_*0*_ correspond to pre bleach imaging. The mobile fraction *mf* represents the fraction of molecules that are free to diffuse. It is estimated by using the first timepoint after bleaching (*I*_*0*_) and the median of the last twenty timepoints (*I*_*f*_) in the following expression:

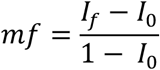

### FLIP Model

Fluorescence loss in photobleaching (FLIP) is used to assess import and export rates of the different constructs. FLIP experiments involve continually bleaching of a region of interest (ROIb) and tracking signal loss from different regions. Quantification of these curves yields the transport dynamics between nucleus and cytoplasm. We set up experiments and analysis motivated from (*20*) for determining the rate of nuclear import and export.

To model the FLIP data, we developed a system of Ordinary Differential Equations (ODEs) describing the change in protein concentration between two compartments i.e., the nucleus and the cytoplasm. These two compartments are linked with boundary fluxes going in (*Q*_*i*_) and out (*Q*_*e*_) of the nucleus (Fig S1).

We assume that the proteins remain in unbound and mobile state in each compartment. During steady state cells maintain a constant ratio (α) for protein concentration between nucleus (*n*) and cytoplasm (*c*), and the flux between both compartments is equal.

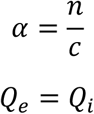

During photobleaching the transport equations for the number of unbleached molecules in nucleus (*N*) and cytoplasm (*C*) can be described as follows, where (*Q*_*b*_) is the number of molecules being bleached per unit time.

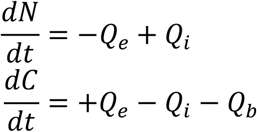

The fluxes are proportional to the concentration of the compartment, times a rate coefficient. Here, *k*_*e*_*’, k*_*i*_*’* are export and import rate coefficients respectively and η’ is the bleaching rate:

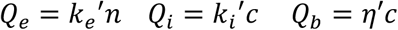

Because these rates (in units of volume per unit time) will depend on the size of the compartment, we define normalized rates as 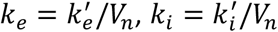, where *V*_*n*_ is the volume of the nucleus. Thus:

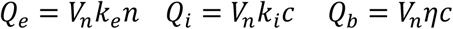

This enables us to rewrite transport equations in terms of concentration.

During bleaching,

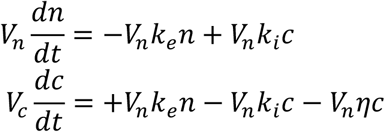

Where *V*_*c*_ is cytoplasm volume. During steady state,

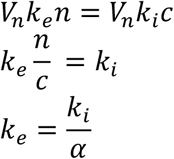

One can further simplify these by using ratio of nuclear volume to cytoplasm volume 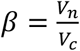

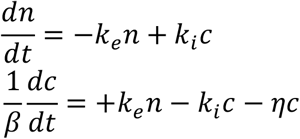

By substituting *k*_*i*_, we get following equations to solve ultimately:

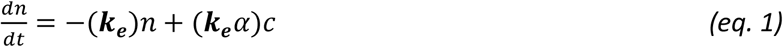

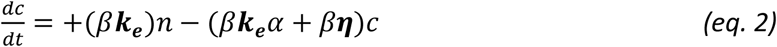

We then solve these equations numerically using MATLAB function ode15s, and fit them to the experimental data to get import/export rates and bleaching rates. Variables in bold are the unknowns to be fitted with fminsearch function in MATLAB (R2020b).

### FLIP Imaging and Analysis

For quantification of FLIP (Fluorescent Loss In Photobleaching) experiments, we followed the fluorescence intensities of three different regions, segmented manually: nucleus, cell, and background. Image acquisition was done with a Zeiss LSM880 inverted confocal microscope using a 63X 1.46 NA oil immersion objective and a 488nm wavelength argon laser. We used a ROI of 17 × 17 (∼12.9 μm^2^) pixels. 10 baseline images were acquired. Then, 40 images of 512 × 512 pixels were acquired every 3 seconds. The power of the laser used to bleach was adjusted to result in the same bleaching rate η. Due to differences in cell morphology, this corresponded to 60% power for cells on 1.5 kPa substrates, and 100% power for cells on 30 kPa substrates. This difference occurred because cells were more rounded on soft gels and therefore thicker in the z axis, leading to a taller column of cytoplasm affected by photobleaching. Cells with beaching rates above 0.12 were discarded. We note that differences in obtained rates between 1.5/30 kPa substrates were reproduced when comparing cells at 30 kPa with/without DN KASH overexpression, where cell morphologies and bleaching laser power was not altered. In the mathematical model, the transport between nucleus and cytoplasm is modelled as transport between two compartments, where the cytoplasm is continuously bleached. We assume that the concentration of protein is uniform in each compartment and that during steady state (before photobleaching) the ratio (α) between nucleus and cytoplasm’s protein concentration is constant. The regions of interest identified for nucleus and cytoplasm were narrow rings around the nucleus, either inside or outside of the nucleus. The average fluorescence intensity of these regions was used as a proxy for nuclear concentration (n) and cytoplasmic concentration (c). The intensities were corrected for background noise, and normalized by the total integrated cell intensity. Experimental data for n and c was used to solve equations 1 and 2, as explained above. The ratio of concentrations at steady state (α) was taken as n/c at the initial timepoint (before photobleaching). To calculate the ratio of nuclear to cytoplasmic volume (β), we first took confocal stacks of cells with labelled nuclei (with DAPI) and whole cell(with GFP), seeded on both 1.5 kPa and 30 kPa gels. In those cells, we noted an excellent correlation between the nuclear/cytoplasmic ratio volume ratio β, and the nuclear/cytosolic area ratio, calculated with nuclear and cytosolic areas at a representative central slice of the cell (Fig. S1). Thus, in FLIP experiments we measured area ratios from images, and converted this to volume rations using the experimental correlation.

To solve for unknown variables, we used a curve fitting technique with a weighted least square method. The experimental data for concentrations (*n,c*) is fitted to a solution of the ODEs (*n*_*f*_, *c*_*f*_). The objective function *f* is then formulated as the sum of squares of residuals of model and experimental data as:

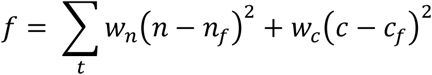

*Where w*_*n*_ and *w*_*c*_ are used to weigh the function by time and compartment concentration to avoid bias in the fitting:

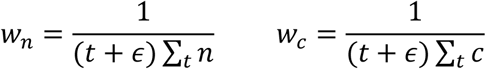

Here, *w*_*n*_, *w*_*c*_, *n, c*, and *n*_*f*_, *c*_*f*_ are all a function of time *t* and *ϵ* is an arbitrary scalar constant (set to 10) used simply to prevent the denominator of *w*_*n*_ and *w*_*c*_ from reaching zero. We use the fminsearch function of MATLAB to minimize *f* as a function of ODE parameters *k*_*e*_ and *η* (equations 1 and 2). For each iteration, *n*_*f*_, *c*_*f*_ is calculated as a function of *k*_*e*_ and *η* using the Matlab ode15s solver.

### Modelling of mechanosensitive nucleocytoplasmic transport

#### Initial conceptual model

To obtain a first understanding of how mechanical force should affect nucleocytoplasmic transport of constructs with NLS sequences, we developed a simple conceptual model. For this, we simply assumed that:

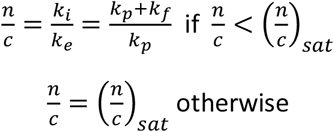

Where n/c is the nuclear to cytoplasmic concentration ratio of a given construct, *k*_*p*_ is a passive diffusion rate through NPCs which decreases with increasing MW (and is equal in the export and import direction), *k*_*f*_ is a facilitated diffusion rate which depends on the strength of the NLS sequence (and does not depend on MW) and 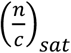 is a maximum, saturated value for n/c ratios. Note that facilitated and passive diffusion are assumed to have additive contributions to total import rates. The effect of force applied to the nucleus is introduced by increasing *k*_*p*_ by two-fold at the lowest MW (arbitrarily set to have to have a value of *k*_*p*_=1 in the absence of force) and by a progressively smaller amount as MW increases, until having a negligible effect at the highest MW (arbitrarily set to have a value of *k*_*p*_=0.015 in the absence of force). Force also increases *k*_*f*_ by 2-fold, in this case independently of MW. After applying these effects of force, mechanosensitivity is calculated as:

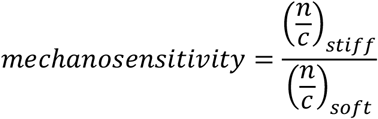

Graphs in fig. 4A,B were calculated by calculating n/c and mechanosensitivity for a range of values of *k*_*p*_ (1-0.015 before force application) and *k*_*f*_, (16-0.12 before force application). The choice of values is arbitrary, and merely intends to show the relative effects when either *k*_*f*_ or *k*_*p*_ dominate the overall n/c ratio. Accordingly, no specific numerical values are shown in the graphs.

#### Kinetic mathematical model of transport

The kinetic model of nucleocytoplasmic transport (Fig. SM1, Tables SM1-SM3) was constructed following a canonical description of the nucleocytoplasmic transport process (*30, 39*–*41*). A system of ordinary differential equations (Table SM1) is used to describe passive diffusion of unbound cargo molecules through NPCs; Ran-mediated facilitated diffusion of cargo:importin complexes through NPCs, and maintenance of the RanGTP gradient across the nuclear envelope through NTF2-mediated import of RanGDP (*69, 70*), RanGAP-mediated hydrolysis of RanGTP to RanGDP in the cytoplasm (*49*), and chromatin-bound RCC1 (RanGEF) mediated conversion of RanGDP to RanGTP in the nucleus (*71*). During passive diffusion, unbound cargo molecules diffuse in either direction at a rate proportional to their concentrations, in accordance with Fick’s law (*24, 72*). During facilitated diffusion, cargo:importin complexes interact with docking sites on NPCs, diffuse across the nuclear envelope and release cargo by interacting with RanGTP. Docking rate to the NPC is proportional to the number of available docking sites. Cargo and importin molecules also associate and dissociate spontaneously in a non-Ran dependent manner.

### Model parametrization

The kinetic model of transport provides a simplified minimal description of the transport process based on a set of canonical assumptions (*30, 39*–*41*). It is not meant to reproduce precise empirical values, rather to characterize dependencies among key biophysical parameters that determine NPC transport kinetics on soft and stiff surfaces. Nonetheless, the model has been carefully parametrized to reproduce key features of transport, and it is remarkably robust to changes in its parameter values. Unless stated otherwise, all simulations were conducted using the mean measured nuclear and cytoplasmic volumes of 627 fL and 2194 fL in our dataset. Passive diffusion rates for different cargo molecules of different sizes were also obtained from measurements (Fig. 1J,K). The cargo concentration was estimated to be in the range 0.01-0.1 *μM*, based on comparison of GFP fluorescence values and reference fluorescence of purified GFP. This is much lower than the ∼10 *μM* physiological concentrations of importins such as Kapß1 (*73, 74*), and the estimated 5-20 *μM* concentration of RanGTP concentration in HeLa cells (*41*), thus precise values of these parameters are expected to have limited effect. Indeed, doubling or halving Ran concentration had limited qualitative effect on our model results. The Ran cycle kinetic parameters were fitted to reproduce a robust nuclear:cytoplasmic RanGTP ratio of >500 (*41*), starting from a 1000:1 ratio. The number of dock sites per NPC was estimated from the thousands of FG binidnig sites per NPC and the large fraction of cargo and NTR molecules found in mass-spectrometry measurements in native NPCs (*75*).

### Simulation code

Our simulations were implemented in Python (version 3.6). They are fully reproducible; the source code and the run parameters can be found in https://github.cs.huji.ac.il/ravehb-lab/npctransport_kinetic/ (run03 was used to produce model results in this manuscript).

**Table SM1.**
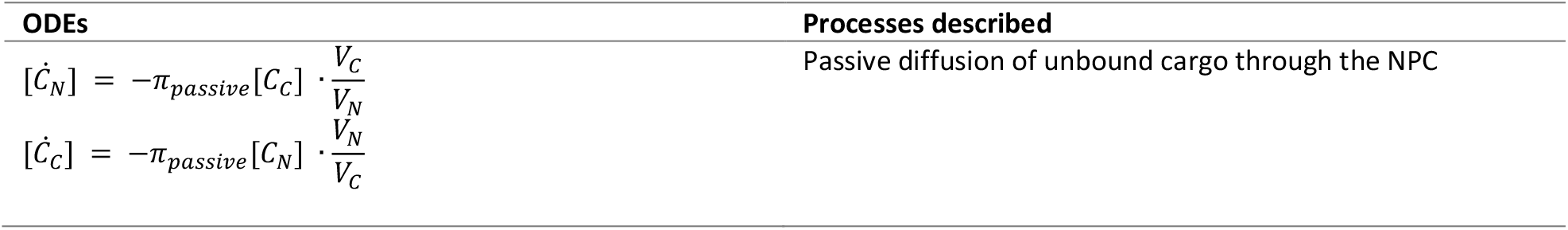
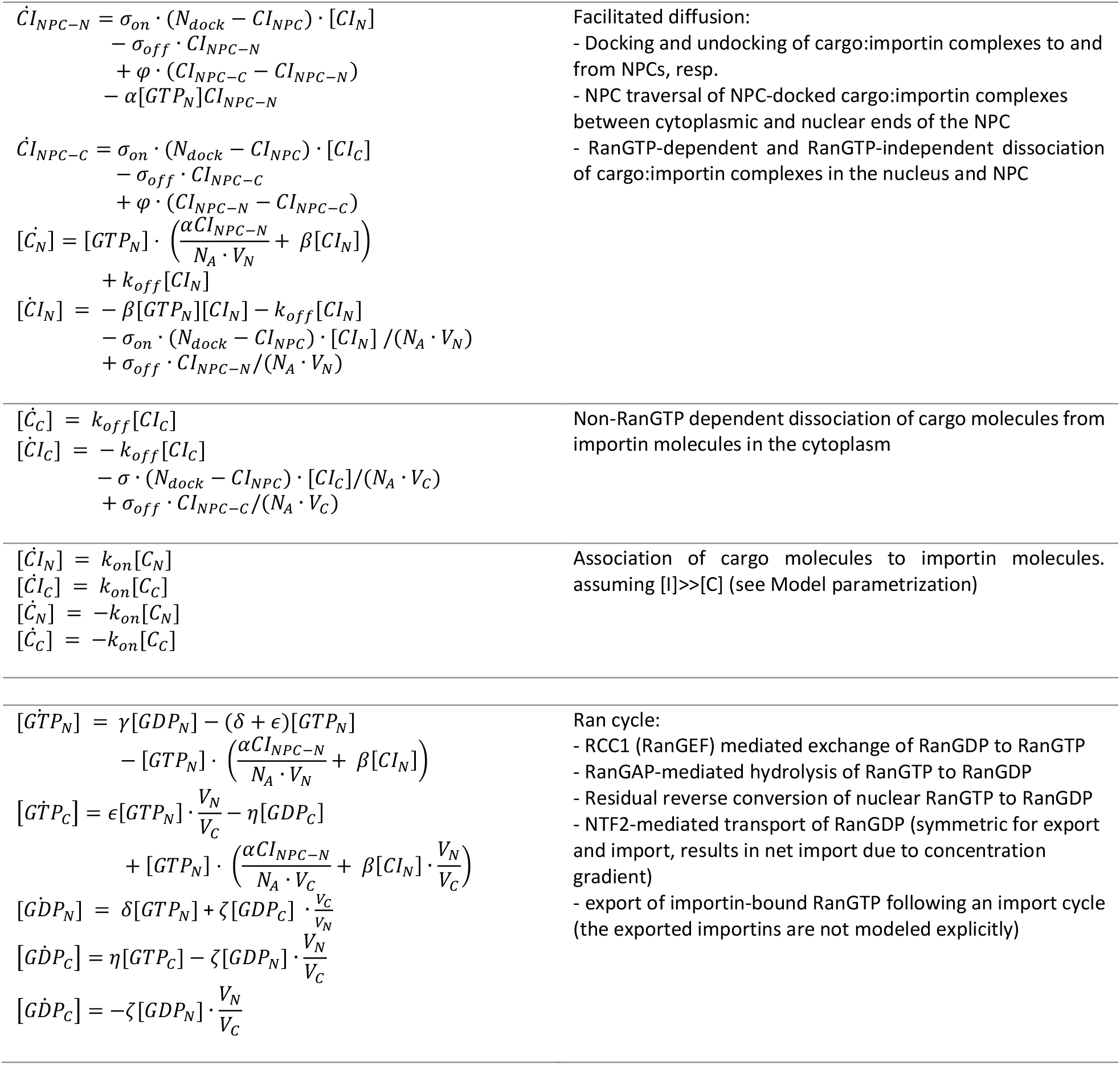
Ordinary differential equations (ODEs) of a kinetic model of transport. Subscripts N and C indicate nuclear and cytoplasmic localization. Subscript NPC indicates localization to the NPC, and subscripts NPC-C and NPC-N indicate sub-localization at the nuclear and cytoplasmic sides of the NPC, respectively. Bracketed variables are in units of concentration (for either the nucleus or the cytoplasm) and non-bracketed variables indicate actual numbers of molecules (for NPC-docked <u>molecules</u>) (Table S1). *N*_*A*_ is Avogadro’s number.

**Table SM2.** Kinetic model variables.

| Variable name | Description |
| --- | --- |
| $C$ | cargo molecules (unbound) |
| $CI$ | cargo:importin complex (bound) |
| $GTP$ | RanGTP |
| $GDP$ | RanGDP |

**Table SM3.**
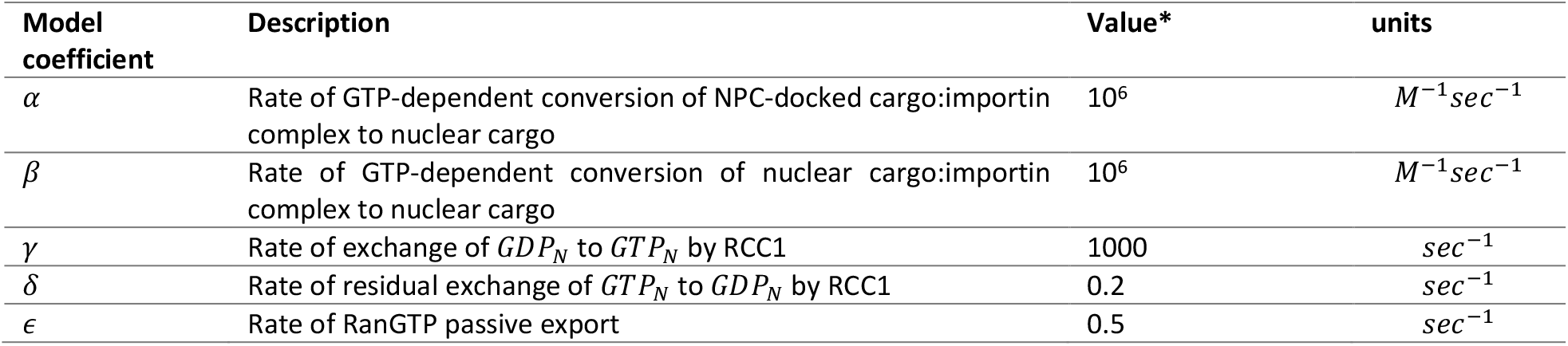
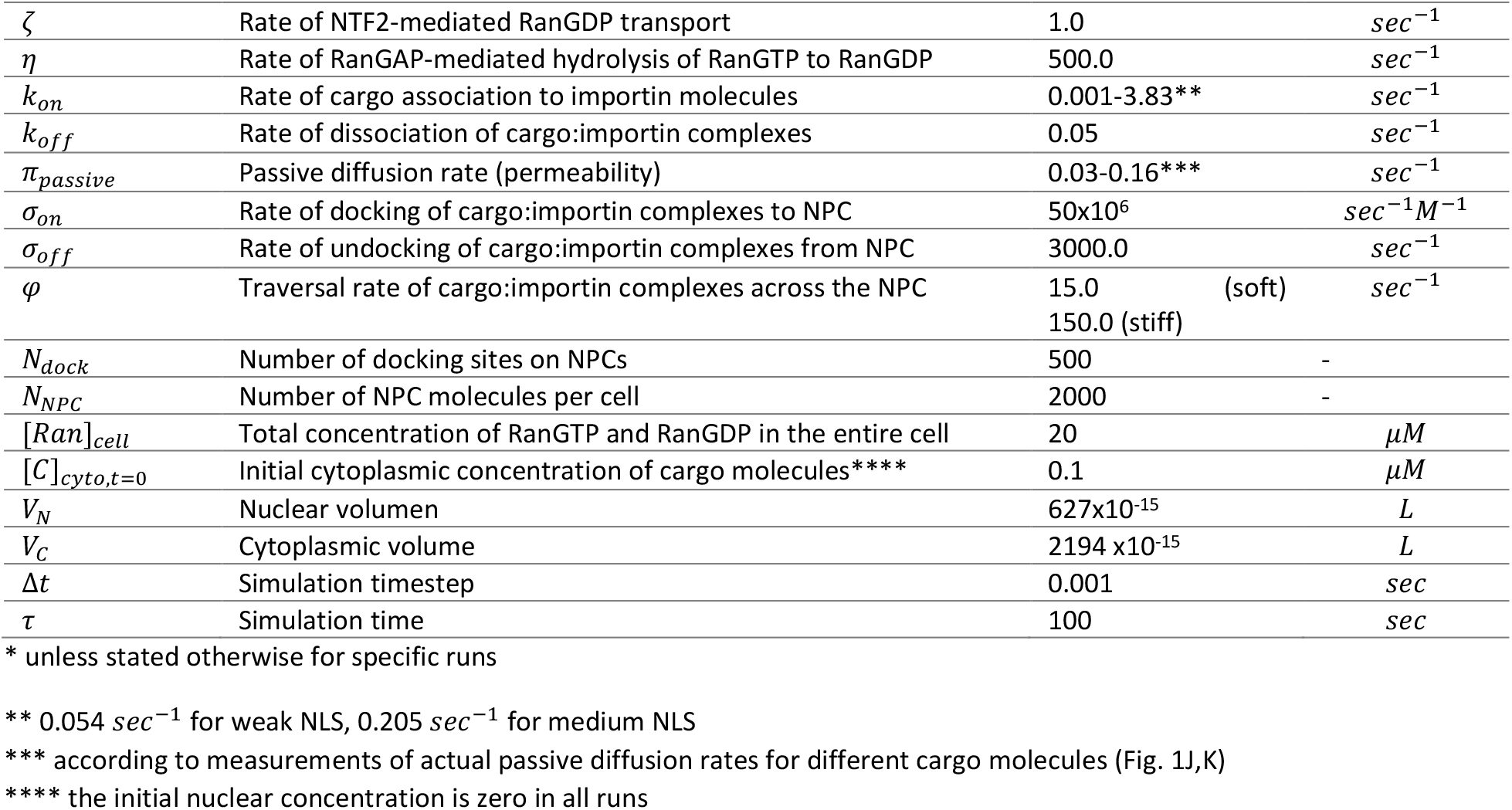
ODE model coefficients.

**Fig. SM1.**
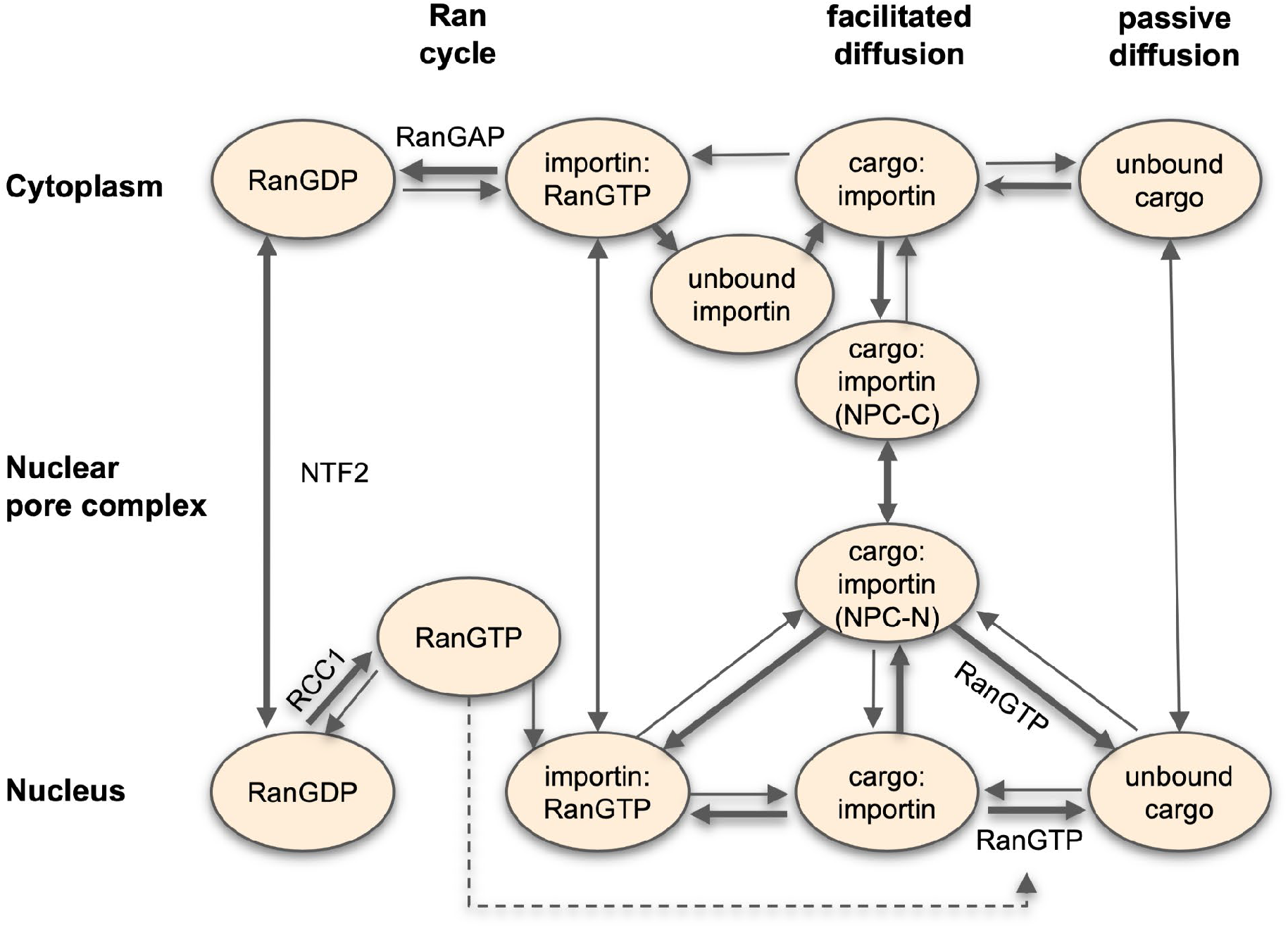
Kinetic model of import through the NPC. The concentration of importin molecules is not modeled explicitly (see Text), except to indicate whether cargo molecules are in the bound or unbound state, but they are shown here for completeness.

### Statistical Analysis

Statistical analysis was performed with GraphPad Prism 9.0.0. When testing data with a 2-way ANOVA, we transformed the data (y=log10(y)) which showed smaller residuals, and therefore better statistical power, when transformed.

## Acknowledgments

We thank S. Usieto for technical support, and JF Abenza, L Rosetti, G Ceada, S. Garcia-Manyes, M. Rout and the members of the Roca-Cusachs and Trepat lab for useful feedback and discussions. We acknowledge funding from the Spanish Ministry of Science and Innovation (PGC2018-099645-B-I00 to X.T., BFU2016-79916-P and PID2019-110298GB-I00 to P. R.-C), the European Commission (H2020-FETPROACT-01-2016-731957), the European Research Council (CoG-616480 to X.T.), the Generalitat de Catalunya (2017-SGR-1602 to X.T. and P.R.-C.), The prize “ICREA Academia” for excellence in research to P.R.-C., Fundació la Marató de TV3 (201936-30-31), and “la Caixa” Foundation (Agreement LCF/PR/HR20/52400004). IBEC is a recipient of a Severo Ochoa Award of Excellence from MINCIN.

## Author contributions

I.A. and P.R.-C. conceived the study; I.A. and P.R-C. designed the experiments; I.A., I.G.-M., M.M.J., A.E.M.B., and A.E.-A. performed the experiments; I.A., I.G.-M., N.R.C., X.T., B.R. and P.R.-C. analyzed the experiments; I.G.-M. designed and cloned all constructs, N.R.C. implemented the FLIP analysis software, K.C. and B.R. implemented the kinetic model of transport, and I.A., I.G.-M., and P.R-C. wrote the manuscript. All authors commented on the manuscript and contributed to it.

---

**Figure S1.**
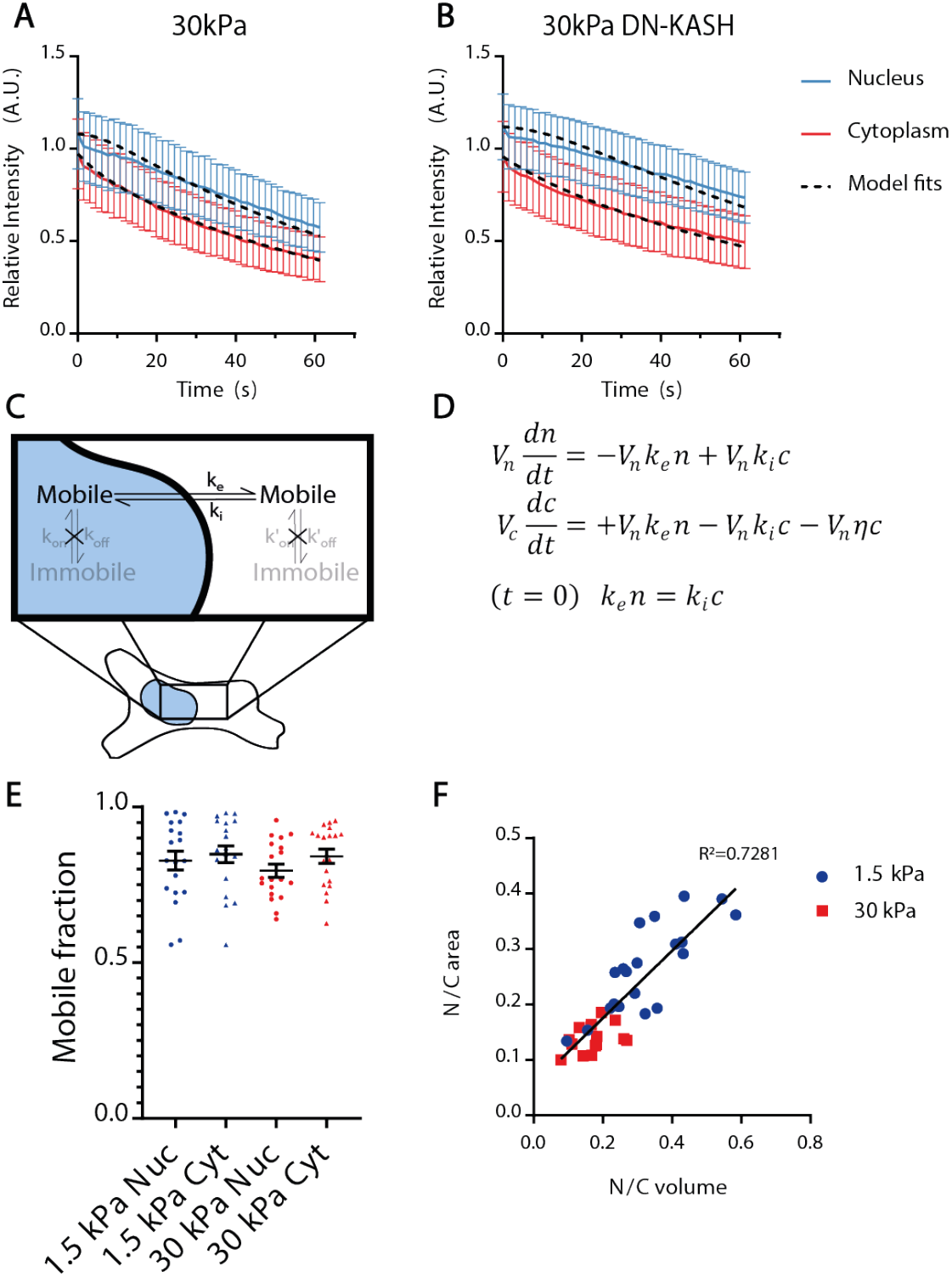
Fluorescence Loss In Photobleaching (FLIP) technique. **A**,**B)** Examples of curves showing fluorescence intensity as a function of time in the nucleus and cytoplasm in FLIP experiments on cells transfected with the diffusive 41kDa construct and seeded on 30 KPa in control condition (A) and with DN-KASH overexpression (B). Data represent the median fluorescence intensity and Standard Deviation of the compartments (nucleus/cytoplasm), normalized with the median of the whole cell before the beginning of photobleaching, and corrected for background signal. Each curve depicts a representative experiment of one cell each. **C**,**D)** Cartoon and equations describing the model used for fitting curves as in A,B, and calculating import and export rates. The model considers the molecules to freely diffuse inside the nuclear and cytoplasmic compartments (see methods). **E**) Mobile fraction of the L_NLS 41kDa construct in the nucleus (Nuc) and cytoplasm (Cyt) of cells seeded on 1.5/30 kPa gels. N=19 cells from 3 independent experiments, lines show mean ±S.E.M. **F**) For cells seeded on 1.5 and 30 kPa gels, correlation between nuclear to cytosolic ratios of volume, and of areas as measured in confocal slices used for FLIP measurements; regression equation *y = 0,6075 x + 0,05375*. N=20 (1.5kPa) and N=14 (30kPa) cells from 2 independent experiments. Black line shows the linear regression.

**Figure S2.**
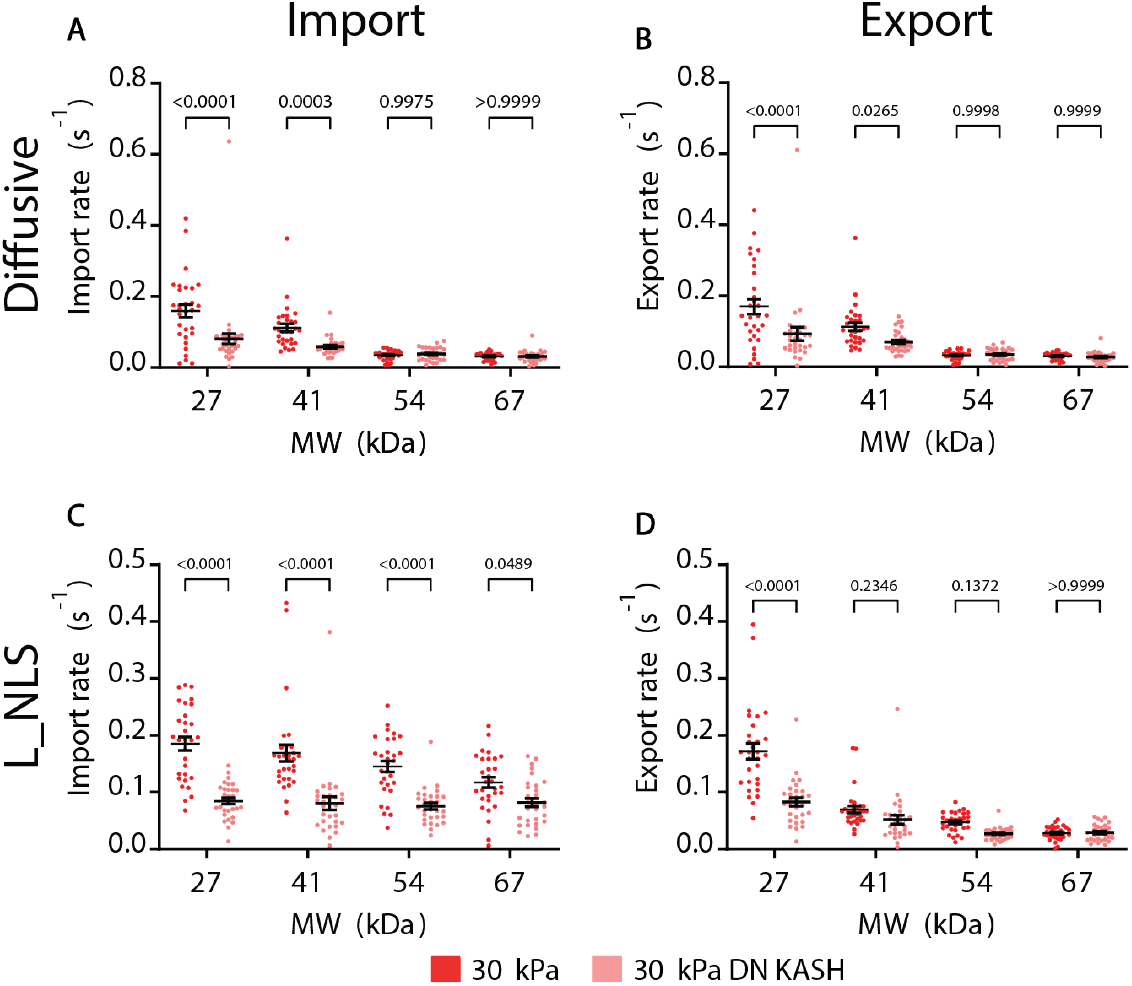
Blocking nuclear to cytoskeletal force transmission with DN-KASH recapitulates the effects of substrate stiffness on transport rates. **A**,**B)** Import and export rates of diffusive constructs for cells seeded on 30 kPa gels, with or without DN-KASH overexpression. In A, both MW (p<0,0001) and Stiffness (p<0,0001) effects tested significant. In B, both MW (p<0,0001) and Stiffness (p=0,0002) effects tested significant. **C**,**D)** Import and export rates of constructs containing L_NLS for cells seeded on 30 kPa gels, with or without DN-KASH overexpression. In C, both MW (p=0,0025) and Stiffness (p<0,0001) effects tested significant. In D, both MW (p<0,0001) and Stiffness (p<0,0001) effects tested significant. N= 30 cells from 3 independent experiments. p-values from Two-way ANOVA. Data are mean ±S.E.M.

**Figure S3.**
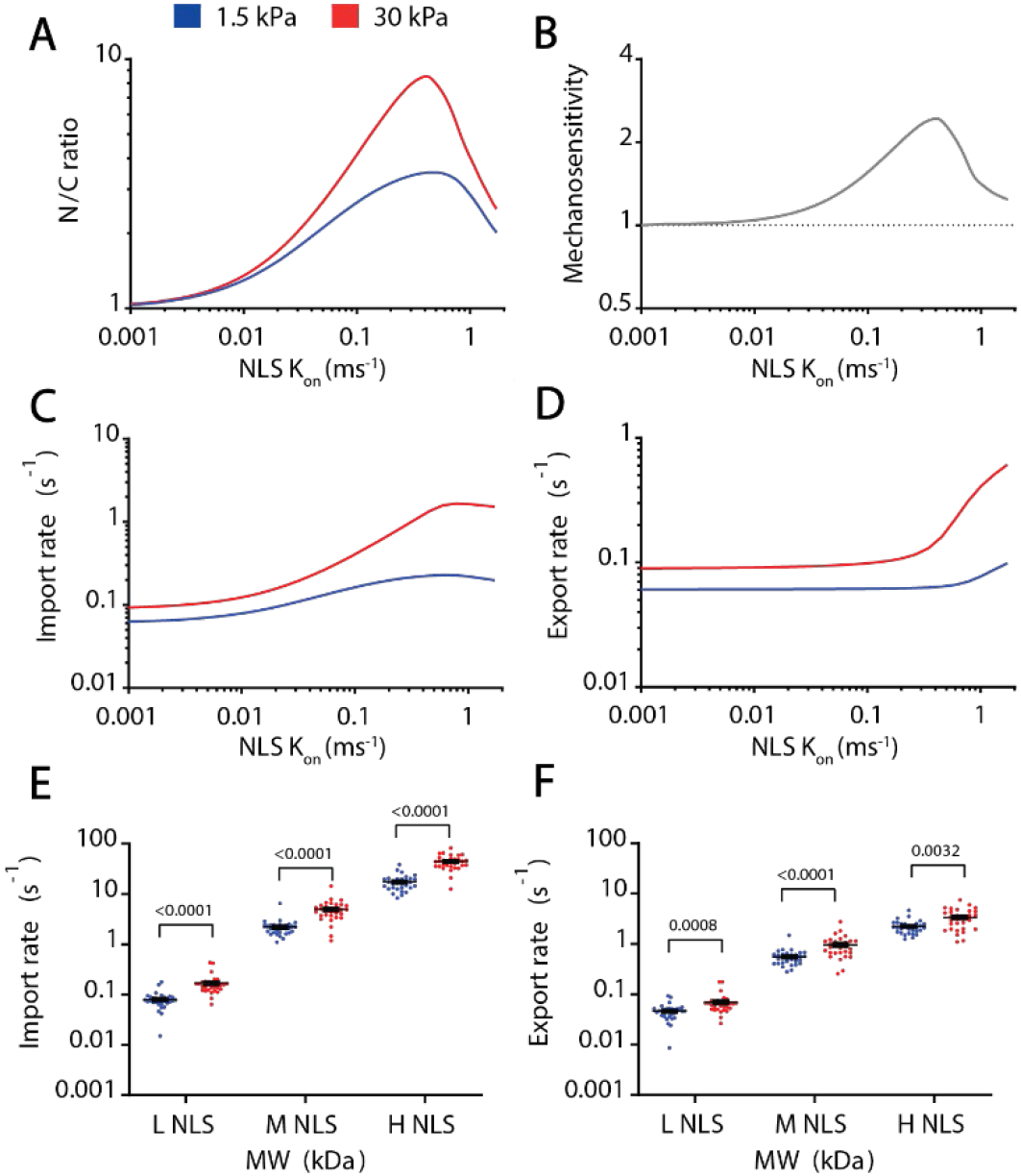
Effect of the affinity of the NLS signal in import and export rates. **(A-D)** Model predictions for N/C ratios (A), mechanosensitivities (B), import rates (C) and export rates (D) for 41kDa constructs as a function of NLS affinity (modelled by the binding rate *kon* between the NLS and importin α). **E-F)** Experimental Import and export rates of 41 kDa constructs containing NLS signals of different affinity for importin β. In both cases (E,F), NLS strength and substrate stiffness effects tested significant (p<0,0001). N= 30 cells from 3 independent experiments. p-values from Two-way ANOVA. Data are mean ±S.E.M.

**Figure S4.**
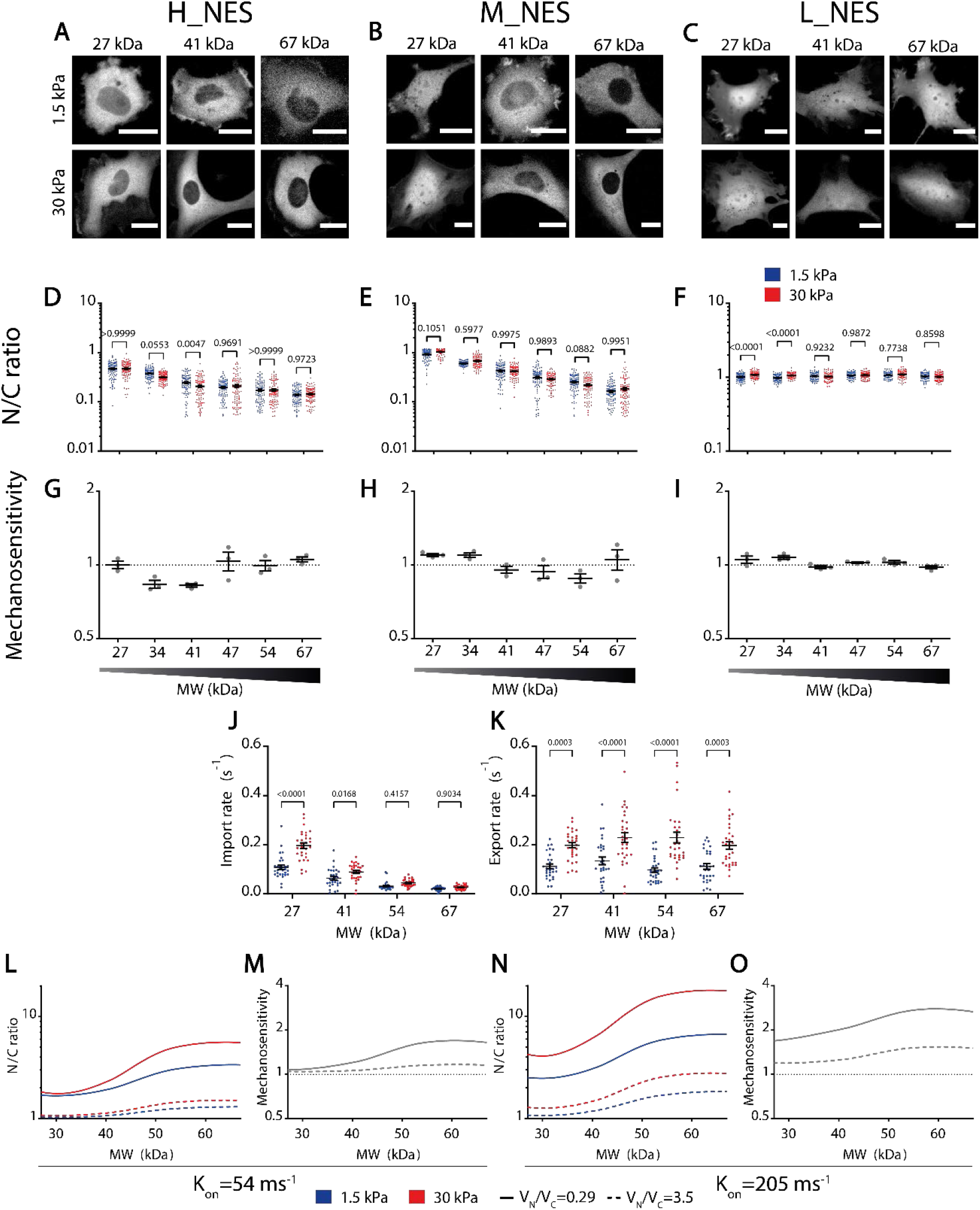
Balance between affinity to Exportin1 and MW defines the mechanosensitivity of nuclear localization in constructs containing NES signals. **A-C)** Representative examples of construct distribution in cells seeded in substrates of 1.5kPa or 30kPa, for H_NES constructs at different MW, M_NES constructs at different MW, and L_NES constructs at different MW. **D-F)** N/C ratios corresponding to the same conditions as A-C. **G-I)** Mechanosensitivity corresponding to the same conditions as A-C. Mechanosensitivity is defined as (N/C)_stiff_/(N/C)_soft_. **J**,**K)** For M_NES constructs, import rates (mediated by passive transport) and export rates (mediated by facilitated transport) as a function of molecular weight. **L-M)** Model predictions of N/C ratios (L) and mechanosensitivities (M) for an NLS with a binding rate k_on_ of 54 ms^-1^ as a function of MW. Data are shown for experimentally measured N/C volume ratios (0.29) and for inverted volume ratios (3.5). **N-O)** Same predictions as in L,M for an NLS with a binding rate k_on_ of 205 ms^-1^. Note that these predictions simply evaluate the role of N/C volumes on import, they do not explicitly model the export cycle (and hence mechanosensitivities are above and not below 1). Statistics: All data were produced in 3 different repeats. D) N= 90 cells from 3 independent experiments. Both MW (p<0,0001) and Stiffness (p=0,0162) effects tested significant. E) N= 120 cells from 3 independent experiments. Only MW effects tested significant (p<0,0001). F) N= 90 cells from 3 independent experiments. Both MW (p<0,0001) and Stiffness (p=0,0001) effects tested significant. Adjusted p-values from 2-way ANOVA; Šídák’s multiple comparisons test. Scale bars: 20 µm. Data are mean ±S.E.M. J,K) Substrate stiffness effects tested significative in both cases (J,K; p<0,0001); MW only tested significative for import (J; p<0,0001).

**Figure S5.**
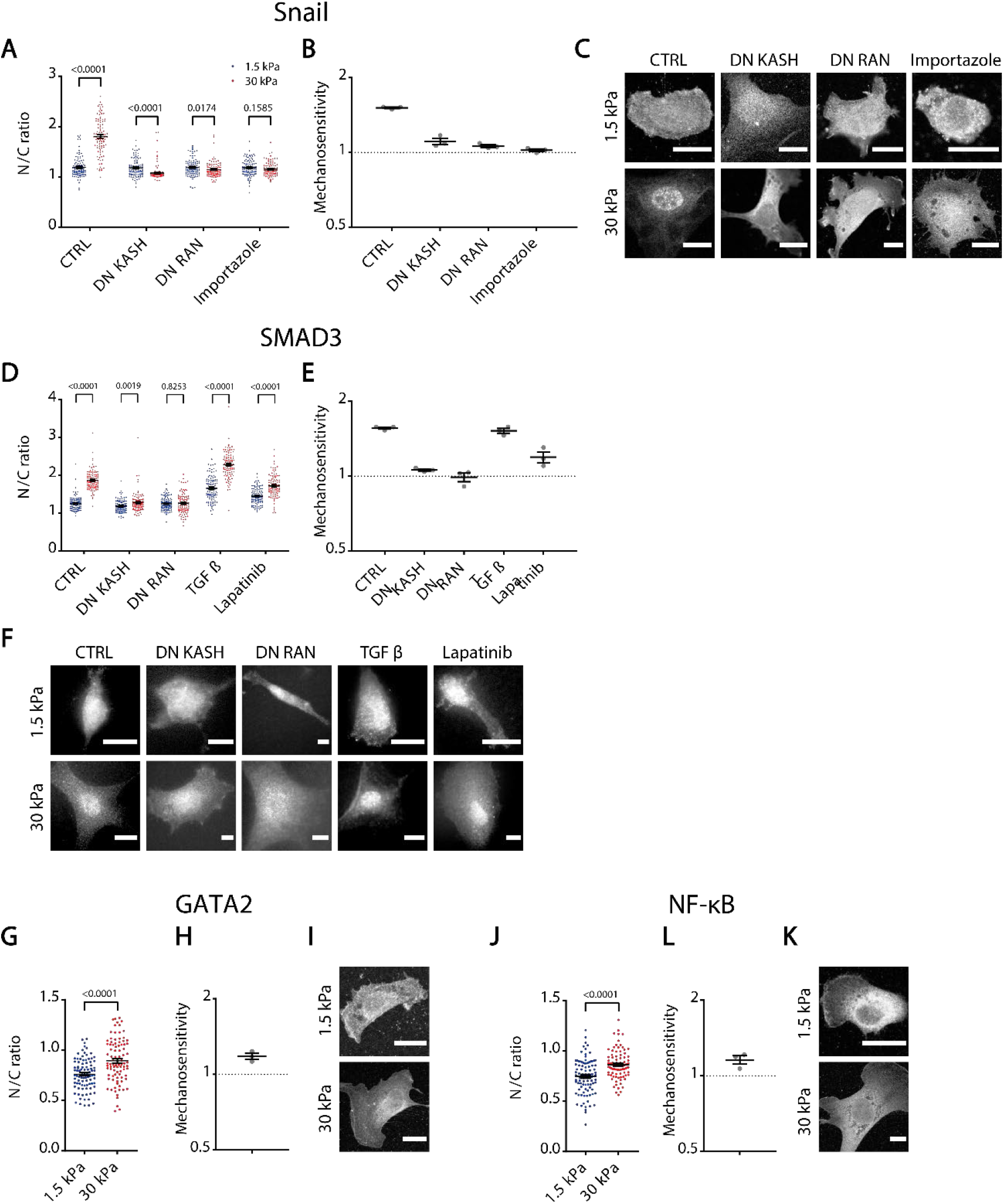
Mechanosensitivity of transcriptional Regulators. **A-C)** For Snail stainings at different conditions, quantifications of N/C ratios on 1.5/30 kPa substrates (A, N= 100 cells from 3 independent repeats), corresponding mechanosensitivities for the 3 different repeats (B), and representative images (C). **D-F)** For SMAD3 stainings at different conditions, quantifications of N/C ratios on 1.5/30 kPa substrates (D, N= 100 cells from 3 different repeats), corresponding mechanosensitivities for the 3 different repeats (E), and representative images (F). **G-I)** For GATA2 stainings at different conditions, quantifications of N/C ratios on 1.5/30 kPa substrates (G, N= 90 cells from 3 independent repeats), Corresponding mechanosensitivities for the 3 different repeats (H), and representative images (I). **J-L)** For NF-κβ stainings at different conditions, quantifications of N/C ratios on 1.5/30 kPa substrates, (J, N= 90 cells from 3 independent repeats), corresponding mechanosensitivities for the 3 different repeats (K), and representative images (L). Scale bars, 20 µm, data are mean ±S.E.M. p-values from corrected multiple Mann-Whitney (A,D) and Mann-Whitney (G,J) tests.

**Table S1.** List of all designed constructs.

| **Name** | **Description** | **Code** |
| --- | --- | --- |
| Diffusive 27kDa (EGFP) | EGFP | IG062/P522 |
| Diffusive 34kDa | EGFP-1PrA | IG024/P277 |
| Diffusive 41kDa | EGFP-2PrA | IG025/P278 |
| Diffusive 47kDa | EGFP-3PrA | IG026/P279 |
| Diffusive 54kDa | EGFP-4PrA | IG027/P280 |
| Diffusive 67kDa | EGFP-6PrA | IG028/P281 |
| L_NLS 27kDa | SV40A4-EGFP | IG065/P525 |
| L_NLS 34kDa | SV40A4-EGFP-1PrA | IG058/P311 |
| L_NLS 41kDa | SV40A4-EGFP-2PrA | IG032/P285 |
| L_NLS 47kDa | SV40A4-EGFP-3PrA | IG059/P312 |
| L_NLS 54kDa | SV40A4-EGFP-4PrA | IG060/P313 |
| L_NLS 67kDa | SV40A4-EGFP-6PrA | IG061/P314 |
| M_NLS 27kDa | SV40A5-EGFP | IG064/P524 |
| M_NLS 34kDa | SV40A5-EGFP-1PrA | IG029/P282 |
| M_NLS 41kDa | SV40A5-EGFP-2PrA | IG031/P284 |
| M_NLS 47kDa | SV40A5-EGFP-3PrA | IG033/P286 |
| M_NLS 54kDa | SV40A5-EGFP-4PrA | IG034/P287 |
| M_NLS 67kDa | SV40A5-EGFP-6PrA | IG044/P297 |
| H_NLS 27kDa | SV40-EGFP | IG063/P523 |
| H_NLS 34kDa | SV40-EGFP-1PrA | IG070/P530 |
| H_NLS 41kDa | SV40-EGFP-2PrA | IG030/P283 |
| H_NLS 47kDa | SV40-EGFP-3PrA | IG071/P531 |
| H_NLS 54kDa | SV40-EGFP-4PrA | IG072/P532 |
| H_NLS 67kDa | SV40-EGFP-6PrA | IG073/P533 |
| L_NES 27kDa | Adeno_NES-EGFP | IG068/P528 |
| L_NES 34kDa | Adeno_NES-EGFP-1PrA | IG046/P299 |
| L_NES 41kDa | Adeno_NES-EGFP-2PrA | IG040/P293 |
| L_NES 47kDa | Adeno_NES-EGFP-3PrA | IG049/P302 |
| L_NES 54kDa | Adeno_NES-EGFP-4PrA | IG050/P303 |
| L_NES 67kDa | Adeno_NES-EGFP-6PrA | IG052/P305 |
| M_NES 27kDa | MAPK_NES-EGFP | IG066/P526 |
| M_NES 34kDa | MAPK_NES-EGFP-1PrA | IG074/P534 |
| M_NES 41kDa | MAPK_NES-EGFP-2PrA | IG038/P291 |
| M_NES 47kDa | MAPK_NES-EGFP-3PrA | IG075/P535 |
| M_NES 54kDa | MAPK_NES-EGFP-4PrA | IG077/P537 |
| M_NES 67kDa | MAPK_NES-EGFP-6PrA | IG051/P304 |
| H_NES 27kDa | HIV_NES-EGFP | IG067/P527 |
| H_NES 34kDa | HIV_NES-EGFP-1PrA | IG045/P298 |
| H_NES 41kDa | HIV_NES-EGFP-2PrA | IG039/P292 |
| H_NES 47kDa | HIV_NES-EGFP-3PrA | IG041/P294 |
| H_NES 54kDa | HIV_NES-EGFP-4PrA | IG042/P295 |
| H_NES 67kDa | HIV_NES-EGFP-6PrA | IG043/P296 |
| Control V5-Twist | V5-Twist | IG106/P641 |
| mut GBP2 V5-Twist | V5-Twist Y107E | IG110/P645 |
| H_NLS-mutNLS V5-Twist | SV40-V5-mTwist K38R K73R | IG115/P669 |
| M_NLS-mutNLS V5-Twist | SV40A5-V5-mTwist K38R K73R | IG116/P670 |
| mutNLS V5-Twist | V5-mTwist K38R K73R | IG117/P677 |
| L_NLS-mutNLS V5-Twist | SV40A4-V5-mTwist K38R K73R | IG118/P678 |
| UL_NLS-mutNLS V5-Twist | SV40A3-V5-mTwist K38R K73R | IG119/P679 |

**Table S2.**
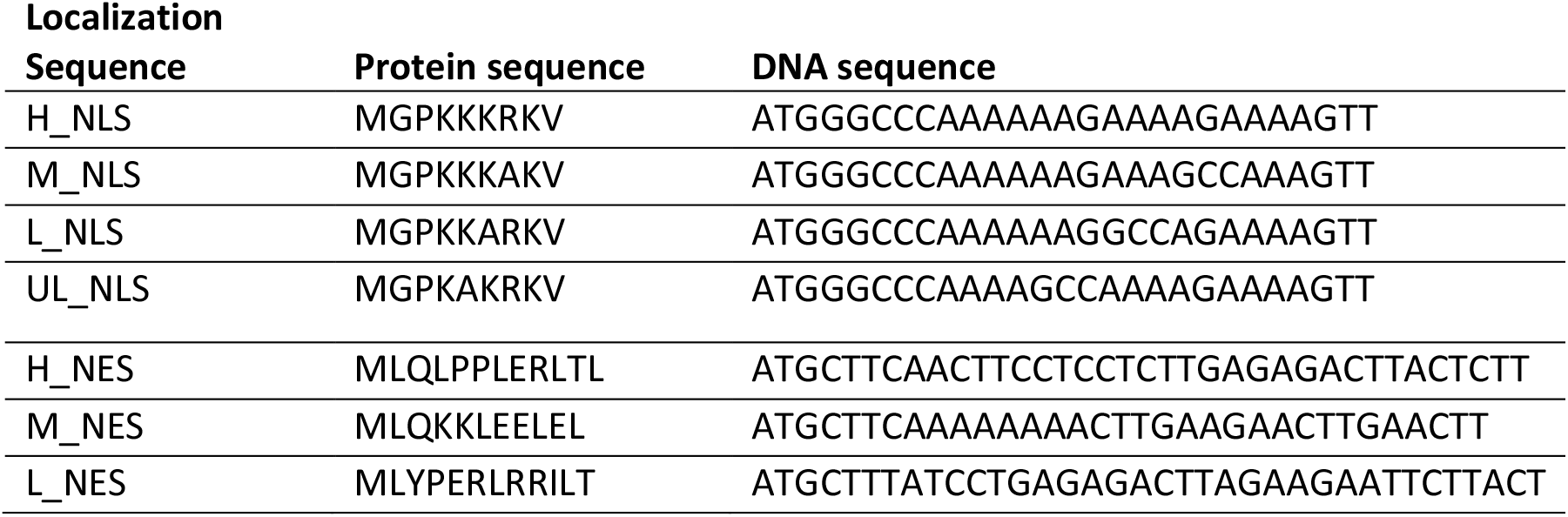
Sequences of NLS and NES sequences used (from refs. (*35*). And (*42*)).

## References

1. F. Broders-Bondon, T. H. N. Ho-Bouldoires, M. E. Fernandez-Sanchez, E. Farge, Mechanotransduction in tumor progression: The dark side of the force. J. Cell Biol. 217 (2018), pp. 1571–1587.

2. O. Hamant, T. E. Saunders, Shaping Organs: Shared Structural Principles across Kingdoms. Annu. Rev. Cell Dev. Biol. 36, 385–410 (2020).

3. J. D. Humphrey, M. A. Schwartz, G. Tellides, D. M. Milewicz, Role of mechanotransduction in vascular biology: focus on thoracic aortic aneurysms and dissections. Circ Res. 116, 1448–1461 (2015).

4. M. L. Lombardi, D. E. Jaalouk, C. M. Shanahan, B. Burke, K. J. Roux, J. Lammerding, The interaction between nesprins and sun proteins at the nuclear envelope is critical for force transmission between the nucleus and cytoskeleton. J. Biol. Chem. 286, 26743–26753 (2011).

5. P. T. Arsenovic, I. Ramachandran, K. Bathula, R. Zhu, J. D. Narang, N. A. Noll, C. a. Lemmon, G. G. Gundersen, D. E. Conway, Nesprin-2G, a Component of the Nuclear LINC Complex, Is Subject to Myosin-Dependent Tension. Biophys. J. 110, 34–43 (2016).

6. A. Elosegui-Artola, I. Andreu, A. E. M. Beedle, A. Lezamiz, M. Uroz, A. J. Kosmalska, R. Oria, J. Z. Kechagia, P. Rico-Lastres, A. L. Le Roux, C. M. Shanahan, X. Trepat, D. Navajas, S. Garcia-Manyes, P. Roca-Cusachs, Force Triggers YAP Nuclear Entry by Regulating Transport across Nuclear Pores. Cell. 171, 1397–1410 e14 (2017).

7. T. J. Kirby, J. Lammerding, Emerging views of the nucleus as a cellular mechanosensor. Nat. Cell Biol. 20, 373–381 (2018).

8. M. M. Nava, Y. A. Miroshnikova, L. C. Biggs, D. B. Whitefield, F. Metge, J. Boucas, H. Vihinen, E. Jokitalo, X. Li, J. M. García Arcos, B. Hoffmann, R. Merkel, C. M. Niessen, K. N. Dahl, S.A. Wickström, Heterochromatin-Driven Nuclear Softening Protects the Genome against Mechanical Stress-Induced Damage. Cell. 181, 800-817.e22 (2020).

9. A. Tajik, Y. Zhang, F. Wei, J. Sun, Q. Jia, W. Zhou, R. Singh, N. Khanna, A. S. Belmont, N. Wang, Transcription upregulation via force-induced direct stretching of chromatin. Nat Mater. 15, 1287–1296 (2016).

10. J. Swift, I. L. Ivanovska, A. Buxboim, T. Harada, P. C. Dingal, J. Pinter, J. D. Pajerowski, K. R. Spinler, J. W. Shin, M. Tewari, F. Rehfeldt, D. W. Speicher, D. E. Discher, Nuclear lamin-A scales with tissue stiffness and enhances matrix-directed differentiation. Science (80-.). 341, 1240104 (2013).

11. A. J. Lomakin, C. J. Cattin, D. Cuvelier, Z. Alraies, M. Molina, G. P. F. Nader, N. Srivastava, P. J. Sáez, J. M. Garcia-Arcos, I. Y. Zhitnyak, A. Bhargava, M. K. Driscoll, E. S. Welf, R. Fiolka, R. J. Petrie, N. S. De Silva, J.M. González-Granado, N. Manel, A. M. Lennon-Duménil, D. J. Müller, M. Piel, The nucleus acts as a ruler tailoring cell responses to spatial constraints. Science (80-.). 370, eaba2894 (2020).

12. V. Venturini, F. Pezzano, F. Català Castro, H.-M. Häkkinen, S. Jiménez-Delgado, M. Colomer-Rosell, M. Marro, Q. Tolosa-Ramon, S. Paz-López, M. A. Valverde, J. Weghuber, P. Loza-Alvarez, M. Krieg, S. Wieser, V. Ruprecht, The nucleus measures shape changes for cellular proprioception to control dynamic cell behavior. Science (80-.). 370, eaba2644 (2020).

13. E. Kassianidou, J. Kalita, R. Y. H. Lim, The role of nucleocytoplasmic transport in mechanotransduction. Exp. Cell Res. 377, 86–93 (2019).

14. X. H. Zhao, C. Laschinger, P. Arora, K. Szaszi, A. Kapus, C. A. McCulloch, Force activates smooth muscle alpha-actin promoter activity through the Rho signaling pathway. J. Cell Sci. 120, 1801–1809 (2007).

15. C. Y. Ho, D. E. Jaalouk, M. K. Vartiainen, J. Lammerding, Lamin A/C and emerin regulate MKL1–SRF activity by modulating actin dynamics (2013), doi:10.1038/nature12105.

16. M. E. Fernandez-Sanchez, S. Barbier, J. Whitehead, G. Bealle, A. Michel, H. Latorre-Ossa, C. Rey, L. Fouassier, A. Claperon, L. Brulle, E. Girard, N. Servant, T. Rio-Frio, H. Marie, S. Lesieur, C. Housset, J.-L. Gennisson, M. Tanter, C. Menager, S. Fre, S. Robine, E. Farge, Mechanical induction of the tumorigenic [bgr]-catenin pathway by tumour growth pressure. Nature. 523, 92–95 (2015).

17. C. Gayrard, C. Bernaudin, T. Dejardin, C. Seiler, N. Borghi, Src-and confinement-dependent FAK activation causes E-cadherin relaxation and beta-catenin activity. J. Cell Biol. (2018), doi:10.1083/jcb.201706013.

18. M. Aragona, A. Sifrim, M. Malfait, Y. Song, J. Van Herck, S. Dekoninck, S. Gargouri, G. Lapouge, B. Swedlund, C. Dubois, P. Baatsen, K. Vints, S. Han, F. Tissir, T. Voet, B. D. Simons, C. Blanpain, Mechanisms of stretch-mediated skin expansion at single-cell resolution. Nature. 584, 268–273 (2020).

19. L. Chang, L. Azzolin, D. Di Biagio, F. Zanconato, G. Battilana, R. Lucon Xiccato, M. Aragona, S. Giulitti, T. Panciera, A. Gandin, G. Sigismondo, J. Krijgsveld, M. Fassan, G. Brusatin, M. Cordenonsi, S. Piccolo, The SWI/SNF complex is a mechanoregulated inhibitor of YAP and TAZ. Nature. 563, 265–269 (2018).

20. N. Ege, A. M. Dowbaj, M. Jiang, M. Howell, S. Hooper, C. Foster, R. P. Jenkins, E. Sahai, Quantitative Analysis Reveals that Actin and Src-Family Kinases Regulate Nuclear YAP1 and Its Export. Cell Syst. 6, 692-708.e13 (2018).

21. E. Jacchetti, R. Nasehi, L. Boeri, V. Parodi, A. Negro, D. Albani, R. Osellame, G. Cerullo, J. F. R. Matas, M. T. Raimondi, The nuclear import of the transcription factor MyoD is reduced in mesenchymal stem cells grown in a 3D micro-engineered niche. Sci. Reports 2021 111. 11, 1–19 (2021).

22. W. SR, R. MP, The nuclear pore complex and nuclear transport. Cold Spring Harb. Perspect. Biol. 2 (2010), doi:10.1101/CSHPERSPECT.A000562.

23. B. M, H. E, The nuclear pore complex: understanding its function through structural insight. Nat. Rev. Mol. Cell Biol. 18, 73–89 (2017).

24. B. L. Timney, B. Raveh, R. Mironska, J. M. Trivedi, S. J. Kim, D. Russel, S. R. Wente, A. Sali, M. P. Rout, Simple rules for passive diffusion through the nuclear pore complex. J. Cell Biol. 215, 57–76 (2016).

25. P. L. Paine, C. M. Feldherr, Nucleocytoplasmic exchange of macromolecules. Exp. Cell Res. 74, 81–98 (1972).

26. D. Mohr, S. Frey, T. Fischer, T. Güttler, D. Görlich, Characterisation of the passive permeability barrier of nuclear pore complexes. EMBO J. 28, 2541–2553 (2009).

27. D. P. Denning, S. S. Patel, V. Uversky, A. L. Fink, M. Rexach, Disorder in the nuclear pore complex: The FG repeat regions of nucleoporins are natively unfolded. Proc. Natl. Acad. Sci. 100, 2450–2455 (2003).

28. M. C. Yuh, G. Blobel, Karyopherins and nuclear import. Curr. Opin. Struct. Biol. 11, 703–715 (2001).

29. M. V. Nachury, K. Weis, The direction of transport through the nuclear pore can be inverted. Proc. Natl. Acad. Sci. 96, 9622–9627 (1999).

30. B. Cautain, R. Hill, N. De Pedro, W. Link, Components and regulation of nuclear transport processes. FEBS J. 282 (2015), pp. 445–462.

31. D. Kalderon, B. L. Roberts, W. D. Richardson, A. E. Smith, A short amino acid sequence able to specify nuclear location. Cell. 39, 499–509 (1984).

32. D. Görlich, Transport into and out of the cell nucleus. EMBO J. 17, 2721–2727 (1998).

33. S. R. Wente, M. P. Rout, The Nuclear Pore Complex and Nuclear Transport. Cold Spring Harb. Perspect. Biol. 2, a000562 (2010).

34. D. Niopek, P. Wehler, J. Roensch, R. Eils, B. Di Ventura, Optogenetic control of nuclear protein export. Nat. Commun. 7, 1–9 (2016).

35. M. R. Hodel, A. H. Corbett, A. E. Hodel, Dissection of a nuclear localization signal. J. Biol. Chem. 276, 1317–1325 (2001).

36. S. K. Lyman, T. Guan, J. Bednenko, H. Wodrich, L. Gerace, Influence of cargo size on Ran and energy requirements for nuclear protein import. J. Cell Biol. 159, 55– 67 (2002).

37. K. Ribbeck, D. Görlich, The permeability barrier of nuclear pore complexes appears to operate via hydrophobic exclusion. EMBO J. 21, 2664–2671 (2002).

38. A. R. Lowe, J. J. Siegel, P. Kalab, M. Siu, K. Weis, J. T. Liphardt, Selectivity mechanism of the nuclear pore complex characterized by single cargo tracking. Nature. 467, 600–603 (2010).

39. S. Kim, M. Elbaum, A Simple Kinetic Model with Explicit Predictions for Nuclear Transport. Biophys. J. 105, 565–569 (2013).

40. T. Jovanovic-Talisman, A. Zilman, Protein Transport by the Nuclear Pore Complex: Simple Biophysics of a Complex Biomachine. Biophys. J. 113 (2017), pp. 6–14.

41. D. Görlich, M. J. Seewald, K. Ribbeck, Characterization of Ran-driven cargo transport and the RanGTPase system by kinetic measurements and computer simulation. EMBO J. 22, 1088–1100 (2003).

42. C. Kanwal, S. Mu, S. E. Kern, C. S. Lim, Bidirectional on/off switch for controlled targeting of proteins to subcellular compartments. J. Control. Release. 98, 379– 393 (2004).

43. S. Dupont, L. Morsut, M. Aragona, E. Enzo, S. Giulitti, M. Cordenonsi, F. Zanconato, J. Le Digabel, M. Forcato, S. Bicciato, N. Elvassore, S. Piccolo, Role of YAP/TAZ in mechanotransduction. Nature. 474, 179–183 (2011).

44. S. C. Wei, L. Fattet, J. H. Tsai, Y. Guo, V. H. Pai, H. E. Majeski, A. C. Chen, R. L. Sah, S. S. Taylor, A. J. Engler, J. Yang, Matrix stiffness drives epithelial-mesenchymal transition and tumour metastasis through a TWIST1-G3BP2 mechanotransduction pathway. Nat. Cell Biol. 17, 678–688 (2015).

45. K. Zhang, W. R. Grither, S. Van Hove, H. Biswas, S. M. Ponik, K. W. Eliceiri, P. J. Keely, G. D. Longmore, Mechanical signals regulate and activate SNAIL1 protein to control the fibrogenic response of cancer-associated fibroblasts. J. Cell Sci. 129, 1989–2002 (2016).

46. T. Furumatsu, T. Kanazawa, Y. Miyake, S. Kubota, M. Takigawa, T. Ozaki, Mechanical stretch increases Smad3-dependent CCN2 expression in inner meniscus cells. J. Orthop. Res. 30, 1738–1745 (2012).

47. A. Mammoto, K. M. Connor, T. Mammoto, C. W. Yung, D. Huh, C. M. Aderman, G. Mostoslavsky, L. E. H. Smith, D. E. Ingber, A mechanosensitive transcriptional mechanism that controls angiogenesis. Nature. 457, 1103–1108 (2009).

48. S. Ishihara, M. Yasuda, I. Harada, T. Mizutani, K. Kawabata, H. Haga, Substrate stiffness regulates temporary NF-κB activation via actomyosin contractions. Exp. Cell Res. 319, 2916–2927 (2013).

49. F. R. Bischoff, C. Klebe, J. Kretschmer, A. Wittinghofer, H. Ponstingl, RanGAP1 induces GTPase activity of nuclear Ras-related Ran. Proc. Natl. Acad. Sci. U. S. A. 91, 2587–2591 (1994).

50. J. F. Soderholm, S. L. Bird, P. Kalab, Y. Sampathkumar, K. Hasegawa, M. Uehara-Bingen, K. Weis, R. Heald, Importazole, a small molecule inhibitor of the transport receptor importin-β. ACS Chem. Biol. 6, 700–708 (2011).

51. H. F, S. Q, L. Y, X. L, X. C, C. F, W. H, L. H, C. Z, L. F, Z. XH, F. XH, H. JJ, L. S, C. YG, HER2/EGFR-AKT Signaling Switches TGFβ from Inhibiting Cell Proliferation to Promoting Cell Migration in Breast Cancer. Cancer Res. 78, 6073–6085 (2018).

52. S. Singh, A. O. Gramolini, Characterization of sequences in human TWIST required for nuclear localization. BMC Cell Biol. 10, 47 (2009).

53. M. Kampmann, G. Blobel, Three-dimensional structure and flexibility of a membrane-coating module of the nuclear pore complex. Nat. Struct. Mol. Biol. 16, 782–788 (2009).

54. S. R. Solmaz, G. Blobel, I. Melcák, Ring cycle for dilating and constricting the nuclear pore. Proc. Natl. Acad. Sci. U. S. A. 110, 5858–5863 (2013).

55. C. B. Wolf, M. Mofrad, Mechanotransduction: Role of nuclear pore mechanics and nucleocytoplasmic transport. Cell. Mechanotransduction Divers. Perspect. from Mol. to Tissues, 417–437 (2013).

56. C. E. Zimmerli, M. Allegretti, V. Rantos, S. K. Goetz, A. Obarska-Kosinska, I. Zagoriy, A. Halavatyi, J. Mahamid, J. Kosinski, M. Beck, bioRxiv, in press, doi:10.1101/2020.07.30.228585.

57. S. Frey, R. Rees, J. Schünemann, S. C. Ng, K. Fünfgeld, T. Huyton, D. Görlich, Surface Properties Determining Passage Rates of Proteins through Nuclear Pores. Cell. 174, 202-217.e9 (2018).

58. E. Infante, A. Stannard, S. J. Board, P. Rico-Lastres, E. Rostkova, A. E. M. Beedle, a. Lezamiz, Y. J. Wang, S. G. Breen, F. Panagaki, V. S. Rajan, C. Shanahan, P. Roca-Cusachs, S. Garcia-Manyes, The mechanical stability of proteins regulates their translocation rate into the cell nucleus. Nat. Phys. 2019 159. 15, 973–981 (2019).

59. I. V. Aramburu, E. A. Lemke, Floppy but not sloppy: Interaction mechanism of FG-nucleoporins and nuclear transport receptors. Semin. Cell Dev. Biol. 68 (2017), pp. 34–41.

60. P. Roca-cusachs, E. Puklin-faucher, N. C. Gauthier, N. Biais, Integrin-dependent force transmission to the extracellular matrix by α-actinin triggers adhesion maturation (2013), doi:10.1073/pnas.1220723110.

61. J. F. Soderholm, S. L. Bird, P. Kalab, Y. Sampathkumar, K. Hasegawa, M. Uehara-Bingen, K. Weis, R. Heald, Importazole, a small molecule inhibitor of the transport receptor importin-β. ACS Chem. Biol. 6, 700–708 (2011).

62. Q. Zhang, J. N. Skepper, F. Yang, J. D. Davies, L. Hegyi, R. G. Roberts, P. L. Weissberg, J. A. Ellis, C. M. Shanahan, Nesprins: a novel family of spectrin-repeat-containing proteins that localize to the nuclear membrane in multiple tissues (2001).

63. N. Kazgan, T. Williams, L. J. Forsberg, J. E. Brenman, Identification of a Nuclear Export Signal in the Catalytic Subunit of AMP-activated Protein Kinase. Mol. Biol. Cell. 21, 3433–3442 (2010).

64. J. Yang, S. A. Mani, J. L. Donaher, S. Ramaswamy, R. A. Itzykson, C. Come, P. Savagner, I. Gitelman, A. Richardson, R. A. Weinberg, Twist, a master regulator of morphogenesis, plays an essential role in tumor metastasis. Cell. 117, 927– 939 (2004).

65. R. Oria, T. Wiegand, J. Escribano, A. Elosegui-Artola, J. J. Uriarte, C. Moreno-Pulido, I. Platzman, P. Delcanale, L. Albertazzi, D. Navajas, X. Trepat, J. M. García-Aznar, E. A. Cavalcanti-Adam, P. Roca-Cusachs, Force loading explains spatial sensing of ligands by cells. Nature. 552, 219–224 (2017).

66. J. N. Lakins, A. R. Chin, V. M. Weaver, Exploring the link between human embryonic stem cell organization and fate using tension-calibrated extracellular matrix functionalized polyacrylamide gels. Methods Mol. Biol. 916, 317–350 (2012).

67. A. Elosegui-Artola, E. Bazellières, M. D. Allen, I. Andreu, R. Oria, R. Sunyer, J. J. Gomm, J. F. Marshall, J. L. Jones, X. Trepat, P. Roca-Cusachs, Rigidity sensing and adaptation through regulation of integrin types. Nat. Mater. 13, 631–637 (2014).

68. M. A. Rapsomaniki, P. Kotsantis, I. E. Symeonidou, N. N. Giakoumakis, S. Taraviras, Z. Lygerou, EasyFRAP: An interactive, easy-to-use tool for qualitative and quantitative analysis of FRAP data. Bioinformatics. 28, 1800–1801 (2012).

69. S. A, B. A, M. IG, Nuclear import of Ran is mediated by the transport factor NTF2. Curr. Biol. 8, 1403–1406 (1998).

70. K. Ribbeck, G. Lipowsky, H. M. Kent, M. Stewart, D. Görlich, NTF2 mediates nuclear import of Ran. EMBO J. 17, 6587–6598 (1998).

71. R. L, K. J, H. A, W. A, Structural basis for guanine nucleotide exchange on Ran by the regulator of chromosome condensation (RCC1). Cell. 105, 245–255 (2001).

72. K. Ribbeck, D. Görlich, Kinetic analysis of translocation through nuclear pore complexes. EMBO J. 20, 1320 (2001).

73. L. E. Kapinos, B. Huang, C. Rencurel, R. Y. H. Lim, Karyopherins regulate nuclear pore complex barrier and transport function. J. Cell Biol. 216, 3609–3624 (2017).

74. A. Paradise, M. K. Levin, G. Korza, J. H. Carson, Significant Proportions of Nuclear Transport Proteins with Reduced Intracellular Mobilities Resolved by Fluorescence Correlation Spectroscopy. J. Mol. Biol. 365, 50–65 (2007).

75. S. J. Kim, J. Fernandez-Martinez, I. Nudelman, Y. Shi, W. Zhang, B. Raveh, T. Herricks, B. D. Slaughter, J. A. Hogan, P. Upla, I. E. Chemmama, R. Pellarin, I. Echeverria, M. Shivaraju, A. S. Chaudhury, J. Wang, R. Williams, J. R. Unruh, C. H. Greenberg, E. Y. Jacobs, Z. Yu, M. J. de la Cruz, R. Mironska, D. L. Stokes, J. D. Aitchison, M. F. Jarrold, J. L. Gerton, S. J. Ludtke, C. W. Akey, B. T. Chait, A. Sali, M. P. Rout, Integrative structure and functional anatomy of a nuclear pore complex. Nat. 2018 5557697. 555, 475–482 (2018).

